# The electrogenicity of the Na^+^/K^+^-ATPase poses challenges for computation in highly active spiking cells

**DOI:** 10.1101/2024.09.24.614486

**Authors:** Liz Weerdmeester, Jan-Hendrik Schleimer, Susanne Schreiber

**Author notes:** Correspondence: Susanne Schreiber < >.

## Abstract

The evolution of the Na^+^/K^+^-ATPase laid the foundation for ion homeostasis and electrical signalling. While not required for restoration of ionic gradients, the electrogenicity of the pump (resulting from its 3:2 stoichiometry) is useful to prevent runaway activity. As we show here, electrogenicity could also come with disadvantageous side effects: (1) an activity-dependent shift in a cell’s baseline firing and (2) interference with computation, disturbing network entrainment when inputs change strongly. We exemplify these generic effects in a mathematical model of the weakly electric fish electrocyte, which spikes at hundreds of Hz and is exposed to abrupt rate changes when producing behaviorally-relevant communication signals. We discuss biophysical strategies that may allow cells to mitigate the consequences of electrogenicity at additional metabolic cost and postulate an interesting role for a voltage-dependence of the Na^+^/K^+^-ATPase. Our work shows that the pump’s electrogenicity can open an additional axis of vulnerability that may play a role in brain disease.

## 3 Introduction

The evolution of P-type ATPases in ancestral methanogenic archaea (1,2) also lay the foundation for the energetics of energy-intensive signaling tissues like the metazoan nervous systems billions of years later (3). In particular, one of the ATPases, the Na^+^/K^+^-pump, plays a significant role in charging the batteries required to operate the nervous systems that the motile life of all multicellular animals is so reliant on.

The Na^+^/K^+^-pump exchanges intracellular sodium for extracellular potassium ions in a 3:2 ratio, and thereby generates a net outward current. This electrogenic property of the pump appears not only as a useful exaptation for osmoregulation in eukaryotes (4), but also invokes activity-dependent changes in nerve cell excitability (5), mediating hyperpolarization after repetitive stimulation and firing-rate adaptation (6–12). These mechanisms are in turn exploited in specific neuronal encoding paradigms (13–18), cell-intrinsic bursting dynamics (16,19–21), and accelerated ion homeostasis (22–24) and thus pose an example of jury rigging in evolution. In the study at hand, we show that for nerve and muscle cells that need to be tonically active for long stretches of time (on the order of minutes to hours), the electrogenicity of the pump can have further, less explored consequences requiring special adaptations of the biophysical components that underlie sustained electrical activity.

Generally, it is assumed that the Na^+^/K^+^-ATPase instantaneously restores ionic gradients, ensuring the robustness of electrical signaling. Furthermore, it is assumed that the net current that is produced by pump activity has limited effects on spiking activity. Under these assumptions, generally, pump currents are not explicitly modeled and reversal potentials are kept constant at all times. At first glance, this seems a reasonable pragmatic approach both for the interpretation of experimental data as well as for computational models of neural dynamics. Accordingly, relatively little attention has been given to the interference between electrogenic pumping and neuronal voltage dynamics, a trend reinforced by the success of models capturing neural dynamics without pump currents, solely based on fixed ion reversal potentials, such as the classical Hodgkin-Huxley model (25).

Here, we demonstrate that, contrary to this notion, electrogenic Na^+^/K^+^-ATPases can exert a significant direct impact on computational properties of highly active excitable cells, and that Na^+^/K^+^-ATPase electrogenicity could pose a challenge for robust spike-based signaling. For strongly active cells operating at high firing rates, additional mechanisms balancing out the pump’s effect on computation may be required, imposing extra costs on cell signaling. We present these effects in a conductance-based computational model of an excitable cell, which was extended to include dynamic ion concentrations and Na^+^/K^+^-ATPases. To isolate the investigated effects of the pump current and support the generalisability of this study to other cell types, the model includes only classic conductance-based sodium and potassium channels. The only active transporter is the Na^+^/K^+^-ATPase, which is responsible for maintaining ionic homeostasis. Modulation of Na^+^/K^+^-ATPase activity occurs via its sensitivity to intracellular sodium and extracellular potassium concentration.

While, due to their generic nature, the described mechanisms may pose challenges for any excitable cell that has to rely on electrogenic pumps, we here showcase them in the electrocyte of the weakly electric fish (26), chosen because of its persistently high firing rates permanently exceeding hundreds of Hz and its, consequently, significant energetic demand. Electrocytes are the cells that make up the electric organ (EO), creating the electric field in the animal’s environment that is vital for communication (27). Electrocyte firing rates, and thus the frequency of the electric organ discharge (EODf), are key for the animal’s survival: the frequency of the resulting oscillating weakly electric field can be sensed by other individuals through electroreceptor afferents (28) and spans a range of 400 Hz across individuals (29). It constitutes the primary signal transmitting information about sex and hierarchy and is also used in interspecific communication. Due the high-frequency spiking activity and thus high ATP requirement of electroyctes (30,31), the EOD cannot only be expected to have been under a severe evolutionary pressure for energetic efficiency, it also exposes the cells to relatively strong electrogenic pump currents that alter cell excitability. The activity-dependent pump currents thus directly influence electrocyte firing rates, as we argue here, complicating the precise regulation of the excitable cells’ activity.

Our model suggests two major effects of electrogenic pumps on computation in highly active excitable cells: (1) a significant shift in a cell’s baseline activity requiring compensation and (2) strong computational side effects of electrogenic pumping in the presence of functionally relevant input changes. While the first effect is intuitive – electrogenic pumping permanently contributes a hyperpolarizing current that requires a compensation to keep firing-rate set points – the second effect is less so. In particular, it can result in unexpected rate changes up to a complete silencing of spiking activity in cases where a drastic firing increase was required and induce spontaneous activity where silencing was required. Even when effects are milder, causing only graded firing-rate changes but not silencing, the pump’s electrogenicity could interfere with cellular computation when spike timing is to be synchronized across excitable cells. As we show, entrainment of electrocytes by their pacemaker neurons can be disrupted in these cases, weakening the EOD and presumably impairing the fitness of weakly electric fish. Based on the identified mechanisms, we suggest a diverse set of biophysical options that may permanently mitigate the side effects of pump electrogenicity in both cases.

To do so, we first isolate the effects of pump currents on cell excitability with constant stimulation. We then demonstrate the impact of pump currents in a simple network context, when an electrocyte is driven by a pacemaker neuron. We outline the consequences of electrogenic pumping for two behaviorally-relevant signals in weakly electric fish: so-called chirps (i.e., short interruptions in the EOD) and frequency rises (transient frequency sweeps). Finally, we discuss how a voltage dependence of the Na^+^/K^+^-ATPase – compared to a pump that adapts its rate exclusively as a function of bulk ionic concentrations and hence only on long timescales – can be useful in alleviating some of the perhaps detrimental effects of electrogenicity on cellular dynamics.

## 4 Results

The electrogenic property of the Na^+^/K^+^-ATPase plays a role in the osmoregulation of single eukaryotic cells (4) as well as, on the organismal level, in marine osmoregulatory organs (32). In metazoan nerve cells, the Na^+^/K^+^-pump restores the cross-membrane ion gradients that generate resting and action potentials, whilst its electrogenicity induces a hyperpolarising membrane current proportional to its pumping activity. The pump rate’s concentration dependence creates a negative feedback loop, leading to an increase in the hyperpolarising current if nerve cells discharge at higher frequencies where more pumping is required (6–12). This feedback loop could thus reduce, terminate or prevent overactivation. On the other hand, for nerve and muscle cells that need to be tonically active for long stretches of time (on the order of minutes to hours), the electrogenicity could pose substantial challenges and its consequences may need to be balanced by additional mechanisms, the latter of which further restrict energetic efficiency, as we demonstrate in the following.

### 4.1 Na^+^/K^+^-ATPase affects tonic spiking

Most computational models of excitable cells follow the principles of conductance-based models for action potential (AP) generation derived by Hodgkin and Huxley (33). Accordingly, they assume that the compensation of the transmembrane ion currents is carried out perfectly so that reversal potentials remain constant and that the hyperpolarizing current introduced by the pump is negligible (25). An explicit inclusion of the pump into mathematical models of such cells is hence not required. To challenge this view for highly active cells, we consider the following scenario: Let’s assume that an electrogenic Na^+^/K^+^-ATPase is added to the Hodgkin-Huxley type spiking mechanism to compensate ion flow across the membrane. Specifically, we illustrate the effects of electrogenic pumps in an experimentally well constrained model of the tonically active electrocyte of weakly electric fish (26) (Fig. 1 A). For simplicity, the Na^+^/K^+^-ATPase in this first case is assumed to only depend on intra- and extracellular ion concentrations as in (19,34), and ATP levels are assumed to be sufficiently high not to impair Na^+^/K^+^-ATPase activity. Specifically, a voltage dependence of the Na^+^/K^+^-ATPase (described in some previous studies, see for example (35–39)), is omitted here, following the approach taken in most previous concentration-dependent models. The layer of complexity that would be added by a voltage dependence of the pump, however, is discussed in section 4.4.

**Figure 1:**
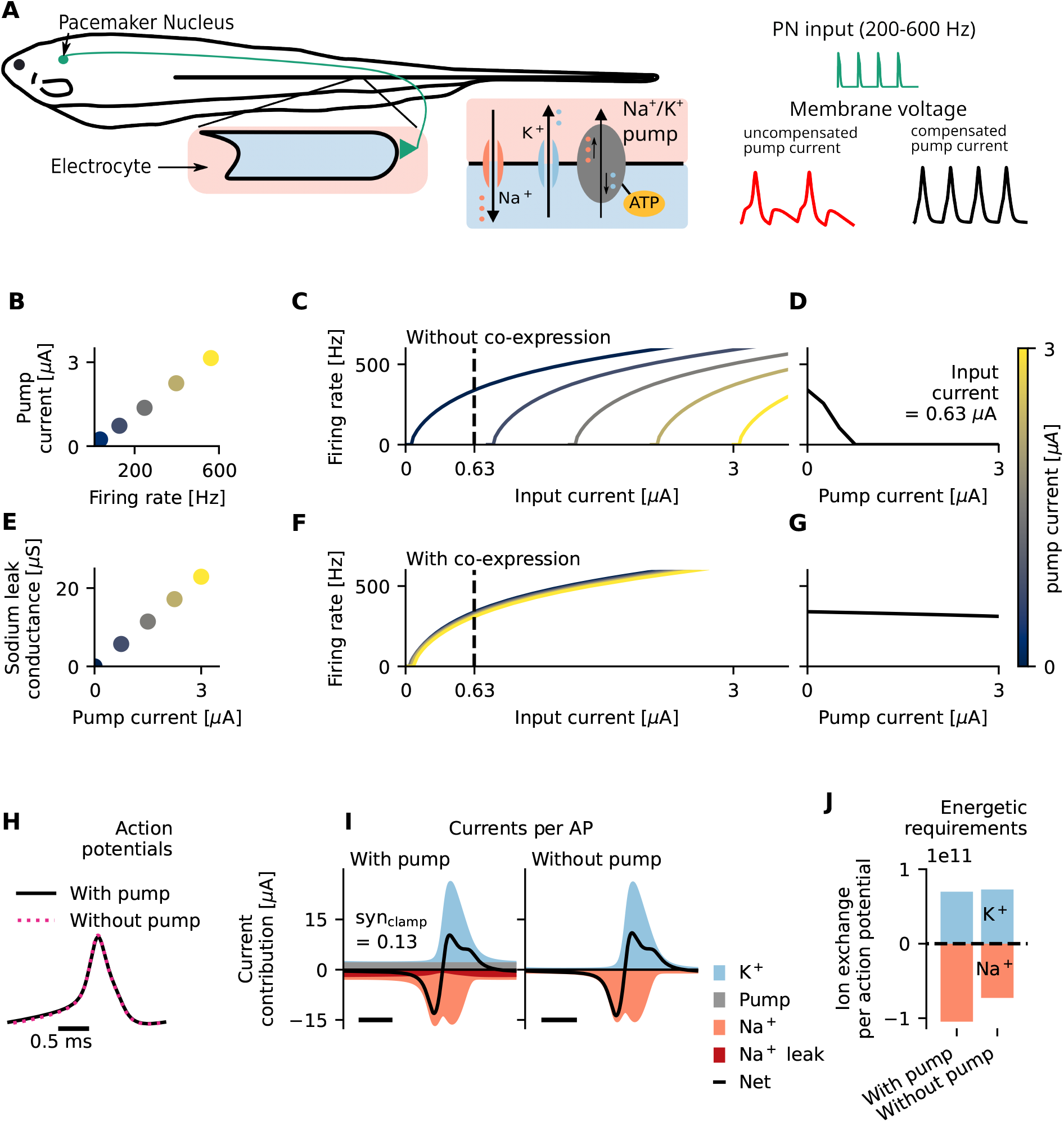
In the weakly electric fish electrocyte, Na^+^/K^+^-ATPase electrogenicity requires compensation, which comes at the cost of a more constrained ion channel composition and sub-optimal energetic efficiency. (A) The weakly electric fish electrocyte is an excitable cell that locks to high-frequency input from an upstream pacemaker (left). Maintenance of ionic homeostasis is carried out by the Na^+^/K^+^-ATPase (grey), which exchanges three intracellular sodium ions for two extracellular potassium ions and thereby generates a net outward current (center). The compensation of this relatively strong pump current is crucial for faithful synchronization to pacemaker inputs (right). (B) High firing rates require significant pump activity, generating a significant hyperpolarising pump current. (C) Increased pump activity, and thus an increased hyperpolarising pump current, reduces cell excitability because a larger inward input current is needed to activate voltage-gated channels. (D) For a fixed physiologically relevant input current (0.63 μA, Methods 6.2.1) that generates tonic firing in an excitable cell, an increase in pump current ‘silences’ the cell. (E-G) Sodium leak channels facilitate a depolarising current that balances out the hyperpolarising pump current. If Na^+^/K^+^-ATPase and sodium leak channels are co-expressed (E), the impact of increased pump activity on cell excitability is minimized (F,G). This is reflected in the impact of the pump current on the tuning curve (F) and the impact of the pump current on the firing rate for a fixed, physiologically relevant input current of 0.63 μA (G). (H-J) Comparison of action potentials and underlying currents for a constant and physiologically relevant synaptic drive (syn_clamp_=0.13, Methods 6.2.1) for a model with and without compensated pump current. (H) Action potentials are similar in size. (I) The additional inward sodium current (dark red) required to balance the outward pump current (grey) results in a simultaneous flow of equally charged ions in opposite directions, decreasing energetic efficiency. (J) Effectively, due to this redundancy more sodium ions per action potential have to be pumped against the gradient.

By measuring sodium currents over time, the pump rates required for ion homeostasis can be estimated((26), Methods 6.1.2). Our model suggests that pump activity sustaining physiological electrocyte firing rates of 200-600 Hz generates a significant hyperpolarising current, here up to 3 *μ*A (Fig. 1 B). Due to the relatively slow dynamics of ion concentrations, on the timescale of the action-potential generation this pump current is approximately constant. Therefore, under the assumption of a constant ion channel composition, a strong hyperpolarising pump current will decrease the input-induced firing rate of the cell, potentially up to the extreme point of silencing it. This effect can be seen in Fig. 1 C, there reflected in a right-shift of the frequency-input curve (tuning curve) of a model with electrogenic pump relative to the model without. In other words, for a constant input that elicits high-frequency tonic spiking without this hyperpolarising pump current, the addition of a pump current decreases a cell’s firing rate; for the strong pump currents that occur in highly active cells (Fig. 1 B), firing is eliminated altogether (Fig. 1 D). To maintain high-frequency firing under physiological pump currents (Fig. 1 B), very high inputs are required (Fig. 1 C). These high inputs could come at a significant metabolic cost of synaptic transmission (specifically the cost related to production and packaging of AChR molecules (26,40)).

Alternatively, the pump-induced raise in current rheobase could be compensated for by adequate adjustments of the cell’s ion channel composition. Specifically, for an excitable cell to operate in a regime of tonic firing, the constant net outward pump current could be balanced by an additional constant inward current, which can be achieved via co-expression of Na^+^/K^+^-ATPases and, for example, sodium leak channels (Fig. 1 E-G, Methods 6.1.1). Although we are not aware of quantitative data on the regulation of ATPase expression in electrocytes, it seems reasonable to assume that the number of pumps expressed in electrocytes scales with the average energetic demand of its spiking activity. An electrocyte that generally fires at higher rates thus requires more pumps to maintain ionic homeostasis. The discharge of the electrical organ (EOD) of *E. virescens* (chosen as a typical representative and whose physiology has been well quantified) approximates to the summed activity of electrocytes. Because individual fish have different baseline EODfs (29), their electrocyte firing rates also differ. Energetic requirements and, consequently, pump expression levels could therefore also vary among individuals. In turn, pump current and the resulting shift in rheobase (that requires compensation) are likely to be unique for each organism. As can be seen in Fig. 1 E-G, an appropriately chosen co-expression factor between Na^+^/K^+^-ATPases and sodium leak channels (Fig. 1 E, Methods 6.1.1) suffices to stabilize the rheobase for a wide range of pump currents (Fig. 1 F-G) which are produced within the regime of physiological firing rates (Fig. 1 B). Therefore, such a co-expression mechanism (similar to those that have been described for homeostatic regulation of intrinsic excitability, see for example (41)) may provide an elegant solution that allows for reliable tonic high-frequency firing with strong pump activity despite cell-to-cell differences in firing rates and pump expression levels.

The proposed co-expression mechanism not only enables reliable high-frequency firing despite electrogenic pump currents, but also compensates pump currents with minimal effects on AP shape (Fig. 1 H). As the overlap of the relatively constant opposing outward pump current with the compensatory inward current, which comprises one-third of the sum of all inward currents (Methods 6.1.2, Eq. 19), results in a largely electroneutral exchange of positive ions (Fig. 1 I), however, a surprisingly high fraction of the pump’s energy is spent on pumping out sodium ions that do not directly contribute to the action potential of the cell but only compensate for the additional pump current. Therefore, the energetic efficiency of action potentials is reduced (42) because of the electrogenicity of the Na^+^/K^+^-ATPase by one third compared to the hypothetical scenario of electroneutral pumping, where no additional inward currents would be needed to enable tonic firing (Fig. 1 J). In absolute terms, this effect is particularly severe for systems operating at high average firing rates which require high pump densities to maintain ionic homeostasis (Fig. 1 B) and even higher pump densities to additionally maintain spike amplitudes (here of around 13 mV) ((26), Appendix 11.1).

In a tonically active cell, the negative feedback loop that the electrogenic Na^+^/K^+^-ATPase provides to enhance ionic homeostasis for action-potential firing could thus come at the cost of a more constrained ion channel composition and sub-optimal energetic efficiency (see Appendix 11.4 for further discussion on metabolic costs).

### 4.2 Na^+^/K^+^-ATPase affects the tuning curve

As outlined above, quantitatively, the compensation required because of the pump’s electrogenicity depends on a cell’s spiking activity. Consequently, even if ion channels and Na^+^/K^+^-ATPases were co-expressed (Methods 6.1.1), and both were optimized to facilitate tonic firing in an excitable cell (Methods 6.1.2), the electrogenicity of the Na^+^/K^+^-ATPase still could interfere with neural processing when firing rates change drastically due to input changes, as outlined in the following.

Most excitable cells do not operate at a fixed firing rate. Often, a flexible, transient modulation of firing rates is required either to track and encode stimuli (43) or to control and adapt motor programs (44). Such a modulation is evoked by alterations in synaptic inputs that differ from baseline activity (Fig. 2 A, D). If the pump rate remains constant despite such changes in firing rates, ion accumulation or drain is to be expected. This can lead to critical transitions in cell excitability and function (as intrinsic cell dynamics depend on ion concentrations (5,21,45,46)), and, in the extreme case, diminish the ion concentration gradient to the extent that firing is completely impaired (47).

**Figure 2:**
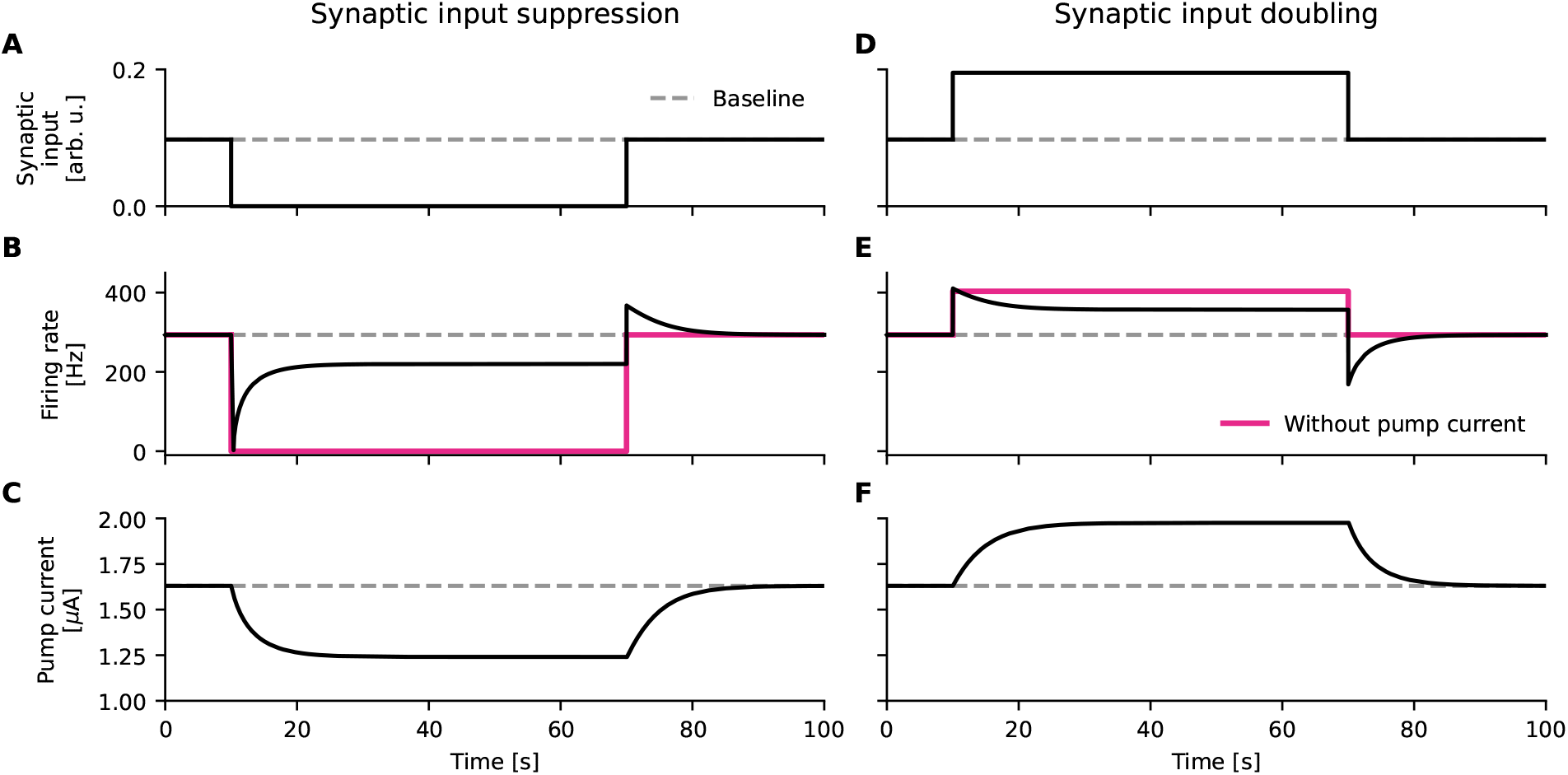
Homeostatic feedback loops based on Na^+^/K^+^-ATPase activity affect firing responses through altered pump currents. (A, B, C) Synaptic input suppression (A) initially silences the cell (B). The Na^+^/K^+^-ATPase adjusts to the reduction in energetic demand through reduced activity. This reduces the pump current (C), which increases cell excitability and results in spontaneous firing without synaptic inputs (B, black). Without a pump current (magenta), spontaneous firing is not induced. (D, E, F) Increased synaptic inputs (D) initially increase firing rates (E). The Na^+^/K^+^-ATPase adjusts to the increase in energetic demand through increased activity. This increases the pump current (F), which decreases cell excitability and results in reduced firing rates (E, black).

At first glance, the pump’s sensitivity to ionic concentrations (Methods 6.1.3) seems an adequate solution that can alleviate such effects of drastically changing firing rates. The dependence of the pump on ionic concentrations contributes to an activity-dependent restoration of ion gradients, i.e., an appropriately calibrated concentration dependence of Na^+^/K^+^-ATPase activity can help to match the energetic demand of the cell’s recent activity (48) (Fig. 2 C, F). The adapted pump activity, however, is accompanied by a change in hyperpolarising current; the cell is pushed to a regime that the system was not originally tuned to (Fig. 2 B, E). Therefore, even in a perfectly controlled environment, an excitable cell could assume different firing rates in response to the same input, where the immediate output of the cell depends on previous activity (16). If a fixed input-output mapping is key to the function of an excitable cell, the electrogenicity of the Na^+^/K^+^-ATPase may induce yet another trade-off between ionic homeostasis and cell function.

### 4.3 Na^+^/K^+^-ATPase affects entrainment

As we illustrate next, a pump-induced alteration of response properties of excitable cells could be especially problematic if these cells are to be entrained in a network or by a pacemaking system. Weakly electric fish electrocytes, like most excitable cells, do not operate in isolation. In order to create an oscillating weakly electric field, electrocyte firing needs to be coordinated across their population, which is enabled by a common drive from an upstream pacemaker. In order to serve a variety of communication paradigms with largely different EOD patterns and hence electrocyte firing rates, an accurate manipulation of the electrocyte firing rates by the pacemaker nucleus is crucial for electric fish.

Each electrocyte in the electric organ is innervated on the posterior side by the spinal motor nerve, which transmits signals from the pacemaker nucleus to the electrocyte (Fig. 1 A) (49). Electrocytes are not synaptically connected among each other; they receive a unidirectional synaptic input from the pacemaker nucleus and firing patterns are only driven by the pacemaker. Therefore, we can model the effects of upstream cells on the electrocytes by simulating the periodic input currents that originate from neurotransmitter release in the synaptic cleft, modulated by pacemaker nucleus activity (Fig. 3 A, top left, Methods 6.2.2) (26). The pacemaker entrains the electrocyte on a spike-by-spike basis, and the electrocyte firing rates should faithfully follow the firing rates of the pacemaker (Fig. 3 A, bottom left) to give rise to a strong, high-amplitude EOD signal.

**Figure 3:**
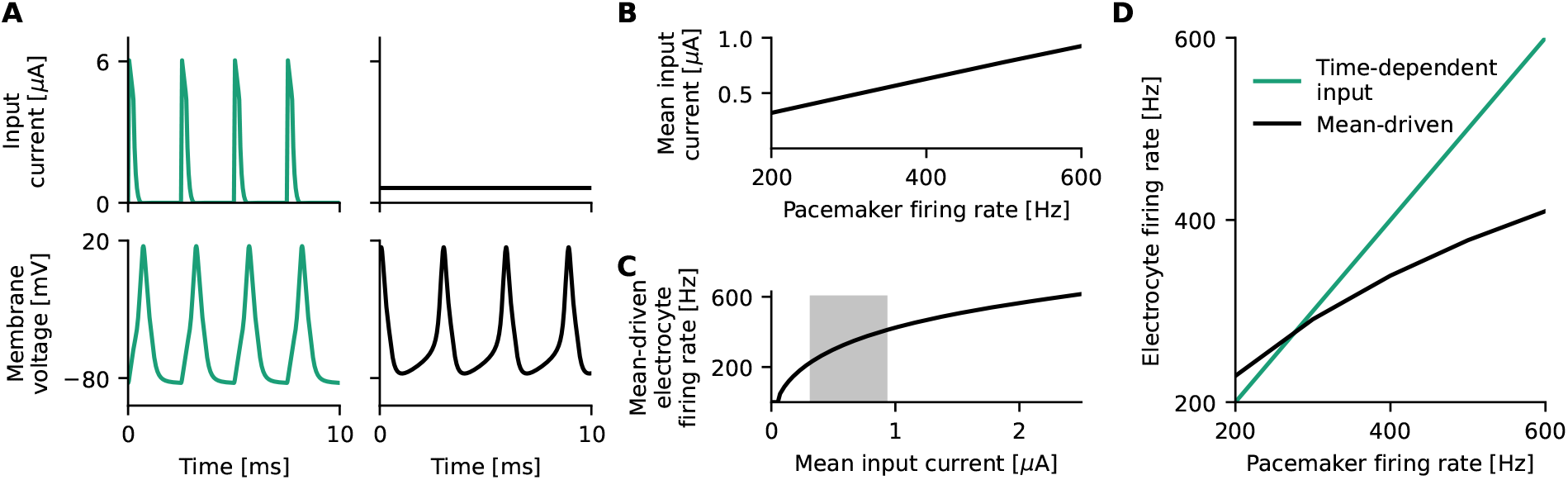
The electrocyte operates in a mean-driven regime, and its mean-driven properties affect its entrainment to periodic inputs from the Pacemaker Nucleus (PN) (A) The input current to the electrocyte stemming from the PN (top, left) sets the high frequency firing rate of the electrocyte (bottom, left). Constant input currents (top, right) also elicit tonic high frequency firing (bottom, right). (B) Mean input currents stemming from physiologically relevant PN inputs of 200-600 Hz. (C) High-frequency electrocyte firing is realized for constant input currents that lie within the mean of the input currents that are generated in the behaviorally-relevant regime (200-600 Hz) (grey box, B). (D) There is a frequency mismatch between the pacemaker firing rate and the mean-driven electrocyte firing rates, which influences signal entrainment.

The model suggests that the electrocyte operates in a mean-driven regime, i.e., the mean of the time-varying input it receives from the pacemaker suffices to invoke tonic spiking in the electrocyte (Fig. 3 A-C). Whether the electrocyte is entrained by the pacemaker depends on characteristics of the electrocyte’s voltage dynamics (like the susceptibility to perturbations, reflected in the so-called phase-response curve) as well as the frequency mismatch between the pacemaker and the firing rate of the electrocyte in response to the stimulus mean. If the frequency mismatch is too large, entrainment fails (Fig. 3 D, Appendix 11.2, Fig. A2). As the pump current affects the mean-driven firing rates of the electrocyte (Fig. 2), it can significantly impact entrainment in this simple network, in particular, when the pacemaker frequency and hence mean input to the electrocyte changes.

For an individual *E. virescens*, pacemaker firing rates can remain constant over long periods of time (50,51). If the electrocyte repeatedly receives the same input and thus produces action potentials that displace a fixed amount of ions per unit of time, pump rates and co-expressed inward leak channels could be tuned to maintain tonic firing and ionic homeostasis (Methods 6.1.2). When searching for food, hiding from predators, and courting, however, substantial deviations from baseline occur. In such cases of drastic change in pacemaker firing rate, the pump rate and thus the pump current adapts via its concentration dependence to the new firing statistics of the electrocyte which, consequently, alters the tuning curve (Fig. 1 C) and hence also the mean-driven firing rates (Fig. 2). In electrocytes of *E. virescens*, behaviorally-relevant deviations from baseline firing come in several forms which include chirps and frequency rises (both used for inter-individual communication), and have different consequences for cell entrainment, as we illustrate in more detail in the following.

#### 4.3.1 The Na^+^/K^+^-ATPase affects the ability to generate EOD chirps

*E. virescens* chirps consist in short cessations (type A) or period doublings (type B) of the EOD, and are thought to play an important role in dominance fights and courtship (29,52,53). Type A chirps, which essentially correspond to short ‘pauses’ in EOD generation, are generated in electrocytes through short interruptions of PN firing (Fig. 4 A, Methods 6.2.2) (54). As electrocytes are only innervated by excitatory synapses, successful chirp generation thus relies on an electrocyte that is ‘silent’ when devoid of input.

**Figure 4:**
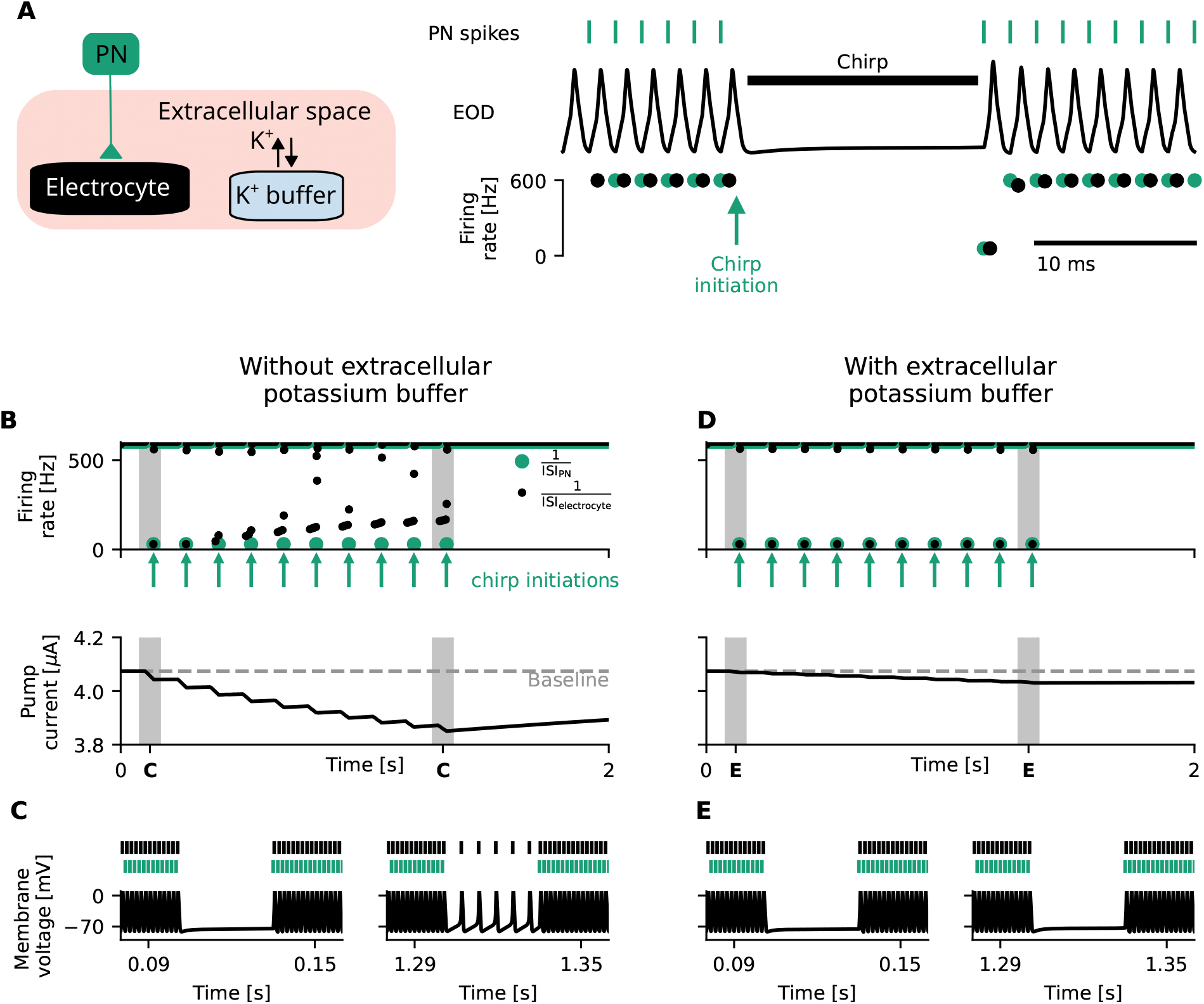
Homeostatic feedback loops on Na^+^/K^+^-ATPase activity impede chirp generation in electrocytes and can be mitigated through extracellular potassium buffering. (A) Schematic illustration of the chirp setting. Left: the electrocyte (black) is coupled to the pacemaker nucleus (PN, green) with an excitatory synapse. A potassium buffer (blue) regulates extracellular potassium concentrations. Right: PN spikes (green, top) induce chirps in the electrocytes through cessation of inputs and thereby temporarily shut off the Electric Organ Discharge (EOD, middle). When chirps are properly generated, instantaneous firing rates (bottom) of the electrocyte (black) equal those of the PN (green). (B) The pacemaker generates ten consecutive chirps, indicated by green arrows and instantaneous PN firing rates 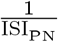. This lowers the mean firing rate of the electrocyte (black, top) and thereby its energetic demand. Consequently, the pump current decreases over time (bottom). This decreased pump current increases cell excitability, which over time (in this paradigm after 400 ms) leads to a mismatch between PN and electrocyte firing rates (top). (C) Electrocyte (black) and PN (green) spikes (top) and electrocyte membrane voltage (bottom) during chirps before (left) and after (right) a significant decrease in the excitability-altering pump current. After such a deviation in pump current, electrocyte firing occurs during chirps (right). (D,E) Same as (B, C) with extracellular potassium buffering. Extracellular potassium buffering extends the timescale of the homeostatic feedback loop of Na^+^/K^+^-ATPase activity on energetic demand which reduces the effect of transient firing-rate deviations on the pump current (D, bottom). This minimizes the deviation in pump current to the extent that chirps can be reliably generated (D (top), E).

From experimental observations it is known that the length of type A chirps in *E. virescens* can extend beyond twenty times the length of one EOD (29,53). Repetitive emission of such long type A chirps (Fig. 4 B, top) decreases mean firing rates of electrocytes and thereby the action-potential-induced ion displacement, ultimately resulting in a lowered pump current (Fig. 4 B, bottom). We find that in our model, the effect of an individual chirp on pump currents is small and does not alter electrocyte excitability to an extent that firing rates are severely affected (Fig. 4 C, left). In case of consecutive chirping, however, the hyperpolarising pump current progressively weakens with time (Fig. 4 B, bottom), eventually leading to spontaneous electrocyte firing in absence of pacemaker input (Fig. 4 C, right). This effect limits the number and duration of chirps that can be induced by the pacemaker (Fig. 4 B, top). The observed dynamics suggest that mechanisms increasing the timescale of the pump feedback loop, such as extracellular potassium buffering (Methods 6.1.4), are suited to diminish the effects of chirps on the pump current (Fig. 4 D) and thereby the effect of variable input signals on chirp generation (Fig. 4 E). We find that extracellular potassium buffering is particularly efficient in dampening Na^+^/K^+^-ATPase effects on cell excitability, because Na^+^/K^+^-ATPase rates of excitable cells are especially sensitive to extracellular potassium concentrations (55,56) (Methods 6.1.3, equation 20) and potassium buffering reduces the variability of potassium concentrations in extracellular space. Metabolic costs of potassium buffering, however, may constitute an additional expense in the total energy budget of the organism (see Appendix 11.4 for further discussion).

#### 4.3.2 The Na^+^/K^+^-ATPase affects generation of frequency rises

During courtship behavior, sudden frequency rises of the EOD followed by an exponential frequency decay back to baseline in the course of 2-40 seconds are thought to constitute an important signal (52). Frequency rises are produced by the pacemaker and the increase in PN firing rate is meant to entrain the electrocyte accordingly (Fig. 5 A). To this end, the mean-driven firing rates of the electrocyte should be sufficiently similar to the transiently elevated PN firing rates, because entrainment fails if frequency mismatches are too large (Fig. 3 D, (57,58)). Accordingly, electrocytes with very slow mean-driven dynamics cannot be entrained to very fast PN inputs (Appendix 11.2, Fig. A2).

**Figure 5:**
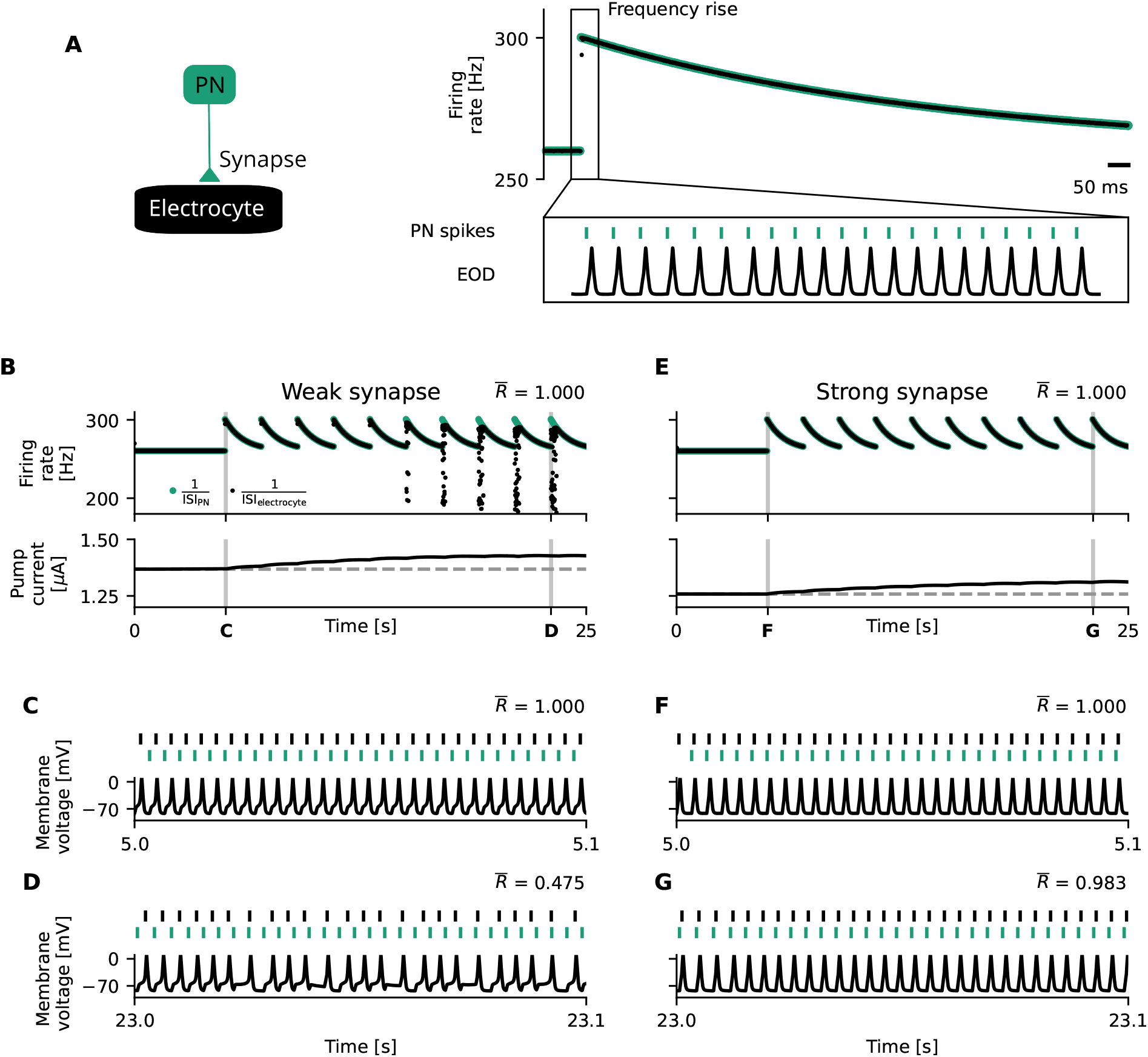
Homeostatic feedback loops on Na^+^/K^+^-ATPase activity impede the generation of frequency rises and can be mitigated through strong synaptic coupling. (A) Schematic illustration of the generation of frequency rises. Left: the electrocyte (black) is coupled to the pacemaker nucleus (PN, green) with an excitatory synapse. Right: Frequency rises are generated through a rapid increase in PN firing rates which exponentially decay back to baseline rates (green, top). As the electrocytes are entrained by the PN (bottom), their firing rates mimic that of the PN and also show a frequency rise (black, top). (B) The generation of consecutive frequency rises by the pacemaker (green) increases the mean firing rate of the electrocyte (black, top) and thereby the energetic demand of the electrocyte, which is fed back into a increased pump current (bottom). This increased pump current decreases cell excitability, which over time (in this paradigm after 15 seconds) leads to a mismatch between PN and electrocyte firing rates during the frequency rises (top). Overall, however, synchronization is very stable, which is reflected in the synchronization index 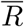 (Methods 6.2.2.1). (C, D) Electrocyte (black) and PN (green) spikes (top) and electrocyte membrane voltage (bottom) during frequency rises before (C) and after (D) a significant increase in excitability-altering pump current. After a significant deviation in pump current, not all PN spikes are reproduced in the electrocyte which leads to ‘missing’ spikes (D). This is reflected in the synchronization index 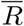, which decreases with increasing pump current deviation. (E-G) Same as (B-D) with strong synaptic coupling. Strong synaptic coupling attenuates the effect of altered pump currents on electrocyte entrainment and enables reliable production of frequency rises (E (top), F, G).

Short frequency rises in *E. virescens* of around 2 seconds have been measured to encompass frequency elevations of up to 40 Hz (29). Repetitive emission of such frequency rises (Fig. 5 B, top) increases mean firing rates and thereby the action-potential-induced ion displacement, resulting in an increased pump current (Fig. 5 B, bottom). Comparable to the observation for chirps, we find that in our model, the effect of a single frequency rise on pump currents is small and does not alter electrocyte excitability to an extent that impedes entrainment (Fig. 5 C). With repetition of these communication signals, however, the hyperpolarising pump current significantly increases over time (Fig. 5 B, bottom), eventually decreasing the electrocyte’s mean-driven firing rate to the point where a precise 1:1 locking between PN and electrocyte breaks down (Fig. 5 D, entrainment index is statistically smaller than in C (59)). Again, this effect of the pump imposes a limit on the number and duration of such frequency rises that can be induced without impairment of electrocyte entrainment and hence the EOD strength (Fig. 5 B, top). Mechanistically, the ability of the electrocyte to entrain to the pacemaker does not only depend on their frequency mismatch, but also on the strength of their synaptic coupling (Appendix 11.2, equation 33). An increase in synaptic coupling strength (Methods 6.1.4) extends the maximum frequency mismatch that still allows for synchronization (Appendix 11.2, equations 38, 37). Both effects of frequency mismatch and synaptic coupling can be illustrated by the so-called Arnold tongue (60). We therefore hypothesize that a strong synapse facilitates electrocyte entrainment and could prolong phases without pump-induced entrainment breakdown (Fig. 5 E-G). Yet it comes at the energetic cost of the increase in neurotransmitter release (including the production and packaging of AChR molecules, see Appendix 11.4 for further discussion) (26,40).

The analysis of both types of signals, chirps and frequency rises, shows that the electrogenic Na^+^/K^+^-ATPase can have significant effects on the computational properties of highly active excitable cells, potentially requiring energetically costly countermeasures for normal operation, especially if cells are to be entrained by a pacemaker or in a network. This suggests that even though Na^+^/K^+^-ATPase was jury rigged to support the generation of action potentials in excitable cells, the fact that their original function required them to be electrogenic, inevitably calls for countermeasures that lower the energetic efficiency of signaling.

### 4.4 The role of Na^+^/K^+^-ATPase voltage dependence

A pump current that only varies on the longer time scale of changes in ion concentrations acts like a constant current on the time scale of spike generation (Fig. 1 I) and will horizontally shift the tuning curves as described above (Fig. 1 C). The consequence of this shift (if uncompensated as described above) is a drastic change in firing rates (Fig. 1 D, Fig. 2, Methods 6.1.1). We next explore whether changes of pump rates on the shorter timescale of action potentials can alleviate the pump-current induced firing-rate adaptation. In particular, we illustrate in the following that a voltage dependence of the pump may constitute an interesting means to limit pump-induced firing-rate modifications and at the same time save on the energetic cost of action potentials.

A common supposition is that (to restore the ionic gradients that get depleted during the generation of action potentials) the pump only depends on ion concentrations and hence, to first approximation, displays constant activity during spiking. In this case, the hyperpolarising pump current counteracts the sodium-currents at the depolarization phase of the AP upstroke and, in fact, assists the potassium currents in repolarising the cell during the action-potential downstroke (Fig. 6 A, left). In the following, we contrast this constant pump to a voltage-dependent pump that activates selectively only during the action-potential downstroke (Fig. 6 A, right). The pump’s voltage dependence could benefit a neuron in two ways: First, by not affecting sodium-based depolarization it may reduce the shift that the pump current induces in the tuning curve. Second, by aiding potassium in repolarization it could provide some energetic benefits for spiking cells.

The exact kinetics and voltage dependence of Na^+^/K^+^-ATPases differ per cell type and organism (38,39,61) and have likely evolved differently in distinct cells to support their unique function and energetic demand. To highlight the positive effects of a voltage dependence of the Na^+^/K^+^-ATPase in electrocytes, whose dependence on voltage to our knowledge currently is unknown, we use a thought experiment based on an idealized voltage-dependent pump that optimally compensates tuning curve shifts and reduces energy demand (Fig. 6 A, right).

Specifically, the dynamics of this idealized voltage-dependent pump is assumed to exactly mimic the hyperpolarising potassium current in the following way: Start with a classical Hodgkin and Huxley equation without pump. Then reduce the repolarising potassium current to 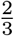 of its strength. The missing 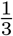 is now substituted by a pump current with exactly the same time course 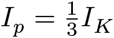, such that,

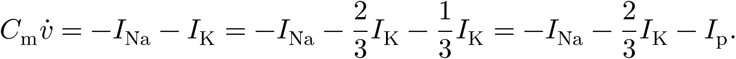

In the model without pump, the cumulative sodium and potassium currents have to add up to zero after one period (Appendix 11.3). This, together with the equation above, implies that the chosen *I*_p_ meets the requirements of the pump stoichiometry 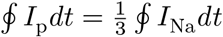 and 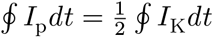 and thereby perfectly counteracts currents flowing during a complete action-potential cycle to maintain ion homeostasis. The equation also shows that replacing one-third of the potassium channels with the idealized voltage-dependent Na^+^/K^+^-ATPase would leave the action potential shape unchanged compared to the model without pump. Importantly, as the pump substitutes for potassium channels, it reduces the flux through these channels by a factor of 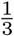. In addition, compared to the constant pump scenario, there is no need for sodium leak channels to cancel out the hyperpolarising pump current. This additionally reduces the cumulative flow of sodium ions by approximately 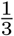 with respect to an excitable cell with relatively constant pumping. Taken together, the reduction in flow of sodium and potassium ions reduces the pump load by approximately 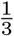 (Fig. 6 B).

Besides lowering the energetic demand, voltage sensitivity can also reduce the effect of the Na^+^/K^+^-ATPases on the neuron’s tuning curve and, consequently, firing-rate adaptation. Two main factors help reducing the adaptation: First, the pump current with a potassium-like voltage dependence is almost completely inactive at spiking onset (approximately −80 mV), in contrast to the fully active constant pump current. Alterations in the former pump current hence induces almost no shift in the tuning curve (Fig. 6 C). Second, as the pump optimally scales with membrane currents at any firing rate in the voltage-dependent case, it instantaneously leads to near-perfect homeostasis. Therefore, action-potential induced concentration changes are minimal. In contrast to the case of a constant pump (Fig. 2), pacemaker-induced jumps in electrocyte drive hence do not elicit a substantial firing-rate adaptation because concentrations remain largely unchanged (Fig. 6 D). We note that even if they would, their effect would be small due to the first argument (Fig. 6 C).

Consequently, suppression of synaptic inputs (like during a chirp) does not result in spontaneous firing for a pump with the described voltage dependence and a cell can be reliably ‘silenced’ for extended periods of time (Fig. 6 D, left). Furthermore, there is a fixed input-output mapping from synaptic input to firing rates in this case (Fig. 6 D, right), which benefits the robustness of entrainment between pacemaker and electrocyte.

While it remains to be established experimentally whether Na^+^/K^+^-ATPases in electrocytes do exhibit a voltage dependence resembling the one postulated here, our thought experiment demonstrates the generic potential of a voltage dependence of pumps to mitigate negative side effects of ion pumping in highly active cells and to lower the need for a costly investment into alternative compensatory measures such as the co-expression of additional ion channels, increased synaptic weights, or extracellular potassium buffering. A voltage dependence of the hypothesized ideal pump, however, does require a higher pump density. As the idealized pump’s activity is constrained to specific time windows, its peak pump rate during these periods needs to be higher. In our model, the effective pump rate during these (limited) times is elevated by a factor of four in comparison to a voltage-agnostic yet constantly active pump (Fig. 6 A). This brings other, potentially significant constraints, such as available membrane space and the cost of pump synthesization and transport to the table (see Appendix 11.4).

## 5. Discussion

We investigated the rarely acknowledged “side effects” of the electrogenic Na^+^/K^+^-ATPase on the computational properties of a highly active spiking cell: the weakly-electric fish electrocyte. Our findings highlight that the electrogenicity of the Na^+^/K^+^-ATPase may pose challenges for robust signal processing in highly active cells; for such cells, a pump that would have evolved for the sole purpose of maintaining ionic gradients would be more efficient if it was electroneutral. We dissect the mechanisms involved and show that energy-intensive countermeasures may be required to ensure robust performance in the presence of the pump when a tonically active cell is driven by inputs of alternating mean – a perspective that is underrepresented in the current literature. Specifically, for the weakly electric fish electrocyte, where robust performance is crucial for interspecific communication, we formulate the following testable hypotheses: *i*) Na^+^/K^+^-ATPase is co-expressed with ion channels that facilitate a relatively constant inward current (such as sodium leak channels). *ii*) In this organism, Na^+^/K^+^-ATPase dynamics have evolved to act in a voltage-dependent manner on the time scale of action potential firing. *iii*) Extracellular potassium is buffered, or Na^+^/K^+^-ATPase sensitivity to this ion is reduced in comparison to that in other organisms. While analysed in electrocytes, the model is and identified mechanisms are sufficiently generic to translate to other excitable cells across the animal kingdom operating at high firing rates, such as Purkinje neurons (62), vestibular nuclear neurons (63), and fast-spiking interneurons (64,65).

When Hodgkin and Huxley established their pioneering conductance-based model of action potential generation in 1952 (25), it was generally assumed that the sodium-potassium pump, which maintains the ionic gradient across the cell membrane, was electroneutral (6). Their computational model, which was adapted over the following 70 years to model numerous types of excitable cells in diverse tissues and species, therefore did not include a pump current. Even later, after the 3:2 stoichiometry of the sodium-potassium pump and thus its electrogenicity were proven, the pump current was often not included in simple point-neuron models, presumably because of its relatively small amplitude and effect size (66,67). Although this may be a suitable argument for excitable cells that are only moderately active, it is less likely to hold for cells that need to be tonically active over long periods of time. In fact, tonically active cells are expected to operate closer to the strong-activity-inducing, posttetanic stimulation protocols used in the 1960s to render to pump current more visible by artificially increasing their effect size (68–70). Additionally, the functional consequences of other, more subtle pump properties, such as its voltage dependence, have not been explored. Here, we identify a range of possible compensation mechanisms for strong pump currents and speculate that, at least theoretically, the pump itself may hold a key to rendering its effects less computationally invasive.

### 5.1 Generalization to other cell types

In this article, two statements were made on the electrogenic pump: In highly active cells its hyperpolarizing current can be strong enough to interfere with signal coordination, and several biophysical cell properties can be exploited to diminish this effect. Assuming that the Na^+^/K^+^-ATPase carries out the majority of active sodium- and potassium transport, on average, the pump current is roughly a third of the sum of all sodium currents (Methods 6.1.2). This holds for any excitable cell (under the above-mentioned assumption), regardless of their ion channel composition, channel biophysics, and additional pumps and transporters. The magnitude of the sum of sodium currents, and thus of the pump current, however, not only depends on a cell’s firing rate, but also on the dynamics of all currents that contribute to the action potential. The time separation between inward sodium and outward potassium currents, for example, dictates AP energy efficiency (42) and thus the pump load. Therefore, excitable cells with different sodium channel dynamics than shown here, such as Purkinje cells with resurgent sodium (71), are likely to exhibit a different degree of crosstalk between the pump current and AP-generating currents, depending on the resulting redundancy in sodium and potassium flows. Moreover, although other ions such as calcium and chloride are assumed not to play a significant role in electrocyte action potentials (27,72), other excitable cells such as mammalian neurons have been shown to be influenced by these ions (73,74).

The main players in calcium removal, which is required to maintain calcium gradients, are the Ca^2+^-ATPase and the Na^+^/Ca^2+^ exchanger (75). Due to its 3:1 stoichiometry, the combined activity of the Na^+^/Ca^2+^ exchanger and the Na^+^/K^+^-ATPase effectively results in an electroneutral active transport of calcium ions. Calcium that is transported through the Ca^2+^-ATPase, however, comes with the byproduct of a strong hyperpolarizing current that will induce the same effects on spiking behavior as the Na^+^/K^+^-ATPase currents presented in this article. Therefore, in cells where calcium currents contribute significantly to its AP generation, the ratio of Ca^2+^-ATPase to Na^+^/Ca^2+^ exchanger expression might be crucial for cell function. The quantative influence of other ions such as calcium and chloride is, however, to be determined in future work.

In general, we can state that as long as sodium ions are the main player in a cell’s AP generation, and if the Na^+^/K^+^-ATPase carries out the majority of active sodium transport, even with additional ionic currents, a highly active cell will experience significant pump currents. We hypothesize that the use of compensatory mechanisms to account for the pump-induced firing-rate adaptation is likely to depend on a cell’s function; if firing-rate adaptation is a desired feature (i.e. in (19)), compensatory mechanisms may not have evolved. In cells such as electrocytes, however, we argue that compensation of pump currents may be required.

Many biophysical mechanisms and cell properties could serve as compensatory mechanisms, of which some have been suggested and modeled in this article. Other possible compensatory mechanisms are discussed below.

### 5.2 Regulatory mechanisms

We discussed four mechanisms that may improve firing-rate control under strong electrogenic Na^+^/K^+^-ATPase currents: co-expression of Na^+^/K^+^-ATPases and sodium leak channels, extracellular potassium buffering, stronger synaptic coupling and pump voltage-dependence. All of these mechanisms can treat the ‘symptoms’ of electrogenic Na^+^/K^+^-ATPase, and could be replaced by any other mechanism that achieves the same effect, i.e., providing an opposing current, diminishing the deviations from baseline pump currents, increasing the entrainment range of a cell, and limiting the pump activity to specific periods of the spike-generation cycle. Some alternative (incomplete list of) compensatory mechanisms that achieve the same effects are discussed below.

Opposing currents do not necessarily have to stem from a sodium leak current, but could also be achieved by other depolarizing currents, such as h-currents (64) or calcium currents (73), or through the (relatively small) current that results from co-transport of H^+^ ions by the Na^+^/K^+^-ATPase itself (76). The former has previously been shown to counterbalance pump currents in the leech central pattern generator neurons (19). Accordingly, the sodium leak also does not have to result from voltage-agnostic channels, but could, for example, be facilitated by a decreased mean half-activation voltage of voltage-dependent sodium channels which could be achieved by transcriptional regulation of channels with different splice forms (77). We lastly note that even though additional supposedly ‘wasteful’ sodium currents might serve a secondary purpose of balancing out fluctuating currents produced via sodium-coupled transport of metabolites (78).

As we showed, deviations in pump currents, resulting from the susceptibility of the Na^+^/K^+^-ATPase to changes in firing rate, can be diminished by prolonging the timescale of the pump’s feedback on cell activity, for example via buffering of extracellular potassium. Similar effects can be obtained from volume increases in intra- and extracellular space or a weaker ion-concentration dependence of the pump. As we showed, for communication with chirps, an uncompensated pump could result in (undesired) spontaneous spiking of electrocytes. In these cases, an increase in the input current required to reach threshold (mediated by additional leak channels) can suppress such activity. Interestingly, compensation could also be provided on the behavioral level: First, an appropriate timing of communication signals, specifically an alternation between frequency rises and chirps, can alleviate undesired changes in baseline pump current due to their opposite impact on pump activity (i.e., increase versus decrease in pump rate). Second, a limitation of individual and cumulative signal duration and amplitude of chirps or frequency rises, respectively, similarly constrains effects on the pump current. This suggests that not only the evolution of ion channels is relevant for shaping communication signals (79) but that also the Na^+^/K^+^-ATPases may have played a significant role.

Finally, we argue that effects of the Na^+^/K^+^-ATPase on neuronal dynamics can be partially avoided if the pump was equipped with an appropriately calibrated voltage dependence. A voltage dependence of the pump has been reported experimentally (35,61). Specifically, the pumping process depends on many individual steps, each involving a different time scale and voltage dependence (36,37). While it remains unclear whether a pump could exhibit a voltage dependence as ideal as in our thought experiment in section 4.4 (in particular with respect to the restriction of its activity to periods of potassium channel activation), at least partial benefits from a modulation of pump activity along some of the qualitative principles described in the thought experiment could be expected. Future experimental investigations on the pump’s voltage dependence, in particular in highly active cells, will therefore be of interest to support or reject the hypothesis predicted from our modeling approach. Interestingly, properties of the pump’s kinetics and voltage dependence have been reported to be highly adaptable in evolution, showing a large heterogeneity in different tissues (38,39), including different splice variants (80) as well as a regulation via RNA editing (61).

### 5.3 Implications for disease

Several neurological diseases (81–84) have been linked to mutations in the *α* subunit of the Na^+^/K^+^-ATPase. The origin of many symptoms observed in these diseases, such as epileptic seizures, lies in pathological neuronal network activity including hyperexcitability and altered oscillatory activity (85). Our modeling work elucidates one mechanism by which altered pump physiology has detrimental effects on cellular computation and can induce pathological network activity. For example, a mutation associated with the rare neurological disease Aternating Hemiplegia of Childhood (AHC) prevents the Na^+^/K^+^-ATPase co-transport of H^+^ ions, the latter of which normally mitigates some of the pump’s negative effects due to its depolarizing contribution (76,86). From our work, we can conclude that the negative side effects of the pump on network computation should be exacerbated by this mutation. We also hypothesize that not only direct impairment of the Na^+^/K^+^-ATPase may contribute to pathological electrical activity, but also deficits in the postulated compensatory mechanisms, thus opening up additional points of physiological vulnerability to pathology.

### 5.4 Other factors of relevance

#### 5.4.1 Reversal potentials

An increase in cellular activity reduces reversal potentials, lowering the ‘driving force’ in action potential generation, and hence affects firing rates (5); vice versa for decreases in activity. The effects of activity-induced concentration changes on neuronal activity therefore stem from a combination of the change in reversal potentials and in pump currents (16). Such effects of the reversal potentials were included in the model and we note that they only contributed mildly (< 5%) to the firing-rate adaptation described.

#### 5.4.2 ATP availability and hypoxia

In addition to ion concentrations, pump rates depend on ATP (87). Limitations in its availability – a phenomenon not uncommon in the weakly electric fish due to its foraging in hypoxic environments (88) – can alter pump rates more drastically than firing-induced changes in ionic concentrations. In theory, a drastic reduction in pump rate (and thus in pump current) results in elevated cell excitability followed by a depolarization block (Fig. 1 C, (21)). In reality, in such cases the EOD of the *E. virescens* only reduces in amplitude but does not significantly change in frequency. The EOD only terminates after very long exposure to annoxia (89), suggesting the existence of compensatory mechanisms diminishing the effects of altered pump currents on cellular activity.

#### 5.4.3 Temperature fluctuations

Weakly electric fish are poikilothermic. Inhabiting affluent streams of the Amazon river (88), with temperature variations of four degrees during a day-night cycle (90), the effects of temperature on spike generation by voltage-dependent channels (91–93) and Na^+^/K^+^-ATPases in *E. virescens* need to be well balanced to keep cell firing in the physiological range.

#### 5.4.4 Spatial effects in electrocytes

Electrocytes are approximately 1 mm long and only excitable on the posterior site (49). The subcellular localization of Na^+^/K^+^-ATPases is hence likely to modulate the impact of the pump’s electrogenicity on cell firing (26). Pumps located on the posterior side are likely to exhibit a more drastic effect on the firing rate of the electrocyte than those on the anterior side. We also note that potential effects of locally-constrained ion concentration changes have, in absence of data about such distributions, been neglected in our model. Constraining changes in ion concentration to specific subcellular locations could influence our results in both directions, either by local amplification or dampening of ion concentration changes (for example via efficient local buffering).

Taken together, our study demonstrates substantial effects of the Na^+^/K^+^-ATPase’s electrogenicity on voltage dynamics in highly active excitable cells. While this property of the most common pump in the nervous system is assumed to serve as a mechanism preventing overexcitability, we show that it can significantly interfere with cellular voltage dynamics via the immediate effects of its highly variable, hyperpolarizing current – posing a particular challenge for highly active cells like electrocytes but perhaps also any other fast-spiking cell in nervous systems. This ultimately calls for strict regulatory mechanisms and may provide an additional evolutionary explanation for the abundance of differential ion channels and the diversity of pump isoforms expressed in excitable cell membranes to not only serve action potential generation, but also the stabilization of firing in biologically realistic environments.

## 6. Methods

All simulations were done in brian2 (94) with a time step of 0.001 ms. We simulated the weakly electric fish electrocyte model from (26), which we incrementally expanded by adding components relevant for modeling the Na^+^/K^+^ pump and corresponding ion concentration dynamics. All parameters were kept the same as in (26), except for the updated and additional parameters (corresponding to additional equations presented in the following section) reported in Table 1.

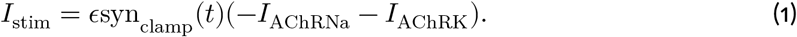

**Table 1:**
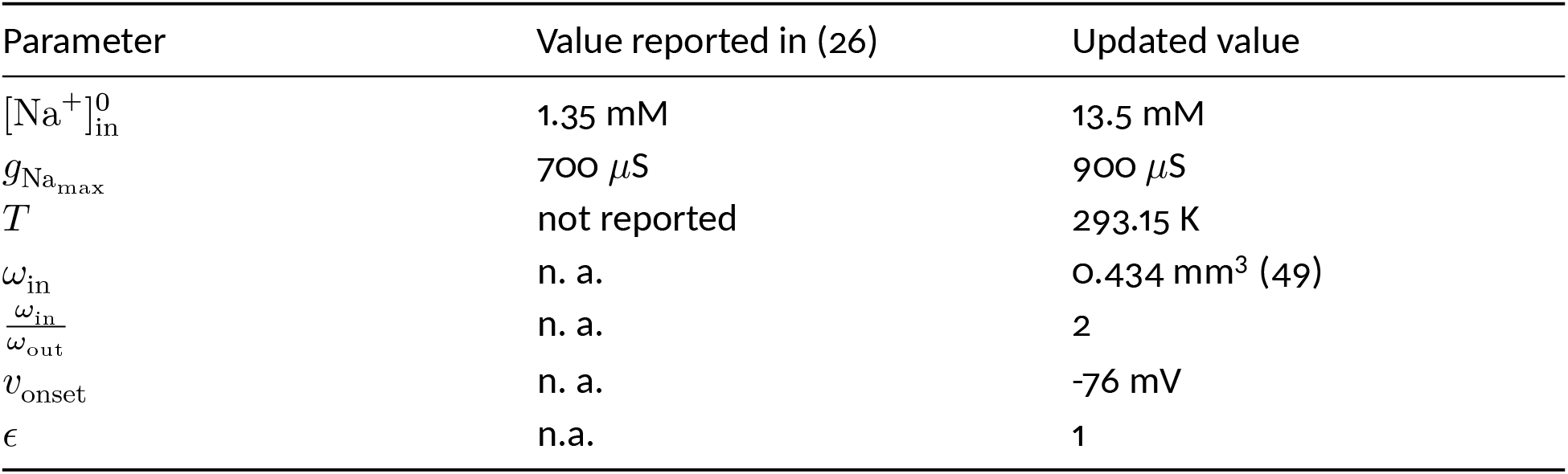
Updated model parameters. The initial (and baseline) intracellular sodium concentration, 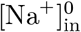, was updated to attain the physiologically relevant sodium reversal potential of 55 mV, and the maximum sodium conductance, 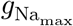 was updated to facilitate a peak amplitude of 13 mV for the mean frequency of the physiological firing range of 200-600 Hz. Model equations were also kept the same as in (26), except for I_stim_ (Eq. 7 in (26)), which was adjusted to correct the directions of current flows through acetylcholine receptor (AChR) channels such that

Note that similarly to (26), the ion flux of calcium ions through AChR channels was neglected under the assumption that similarly to the weakly electric electrogenic wave-type *Sternopygus macrurus* electrocyte, calcium currents do not significantly contribute to action potentials (72).

Furthermore, the synapse strength, *ϵ*, was added as a separate parameter. The additional equations used to model ion concentrations and their derivations are described below in more detail. Note that these are generic, and could be used to expand any point model of an excitable cell where all currents can be separated into sodium and potassium currents.

### 6.1 Modeling an excitable cell with Na^+^/K^+^ pump

First, the Na^+^/K^+^-pump current was added and a suitable co-expression factor between pump density and sodium leak channels was determined in order to counteract the depolarising effects of the pump current. Then, dynamical equations for the ion concentrations were added and the energetic demand at the baseline firing regime was estimated to tune pump densities to maintain steady state ion concentrations. Lastly, the feedback loop of pump density on ion concentrations was modeled to maintain ion homeostasis in firing regimes that deviate from baseline. Each of these steps is explained below in more detail.

#### 6.1.1 Modeling the pump current and sodium leak channel co-expression

In previous work on the computational model of the weakly electric fish electrocyte, it was mentioned that there was an intention to include a Na^+^/K^+^ pump (26). This was however abstained from as the writer noted that adding an additional current *I*_pump_ would leave the model inexcitable. We identified the effects of this pump current on electrocyte excitability, and propose additional currents to counteract these effects.

The Na^+^/K^+^ pump uses one ATP molecule to exchange three intracellular Na^+^ ions for two extracellular K^+^ ions (95). This leads to a net outflux of one positive ion every time the Na^+^/K^+^ pump performs an ion exchange. This net outflux is modeled as an additional current term in the membrane potential evolution equation,

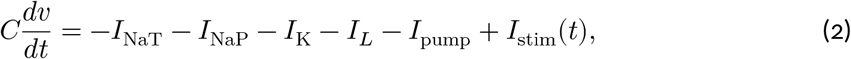

which is the ‘master equation’ that is used for all simulations shown in this study. In the present model, the Na^+^/K^+^ pump strength and thus *I*_pump_ does not depend on the membrane voltage, but on the intra- and extracellular ion concentrations (56). As ion concentration dynamics evolve on much longer timescales than the membrane potential, *I*_pump_ can in this case be assumed to be approximately constant on the timescale of the membrane potential.

When creating the firing-rate (*r*) *vs*. input 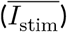 curve (*f*-*I* curve) of the electrocyte, where 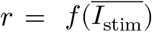, an additional outward pump current creates a horizontal translation of *r*, as 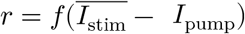. In other words, more inward 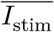 is needed to balance out the outward pump current and push the electrocyte to a firing regime.

To counterbalance the hyperpolarising outward current of the electrogenic Na^+^/K^+^ pump, we introduce an additional inward current. As the first approximation of the Na^+^/K^+^-ATPase current is constant, we modeled this balancing inward current as a relatively constant leak current

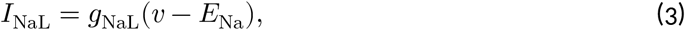

and redefined the leak term in Eq. 2 as

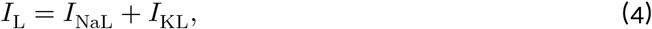

where *I*_KL_ represents the outward potassium leak;

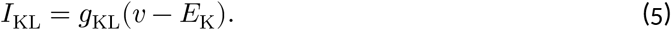

As the original model in (26) has a leak term with a reversal potential equal to *E*_*K*_ (and thus presumably only represents potassium ion flux), *I*_KL_ is the same as the leak term in (26), with the same maximal conductance, *g*_KL_.

For *I*_NaL_ to balance out *I*_pump_, *I*_NaL_ = −*I*_pump_ should hold for all *t*. In contrast to *I*_pump_ however, *I*_NaL_ is highly varying over time as it is dependent on *v*. One condition we can satisfy is for *I*_NaL_ to cancel out *I*_pump_ close to the onset of the *f*-*I* curve. This should render the firing onset of the electrocyte unchanged. We furthermore assume that *g*_NaL_ is adjusted at a larger timescale than *I*_pump_, and that *g*_NaL_ is expressed to counteract *I*_pump_ at a baseline level 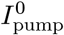. Setting 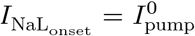 and rearranging to get *g*_NaL_ as a function of 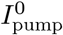 gives

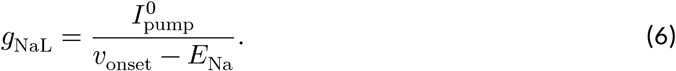

Here, *v*_onset_ is the membrane voltage of the electrocyte just before firing onset. Upon injecting a stimulus of 47nA, we find that *v*_onset_=-81mV. After tuning *g*_NaL_ to counteract the effect of 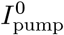 using equation 6, we find that the *f*-*I* curve is least affected by 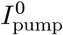 when slightly increasing *v*_onset_ to −76mV. The properties of sodium leak channel co-expression are exemplified in Fig. 1 E-G and implemented to compensate for baseline pump currents in figures 1 B, H-J(with pump), 2, 4, 5, 6 A-C (constant pump), Fig. A1 A1 (with co-expression), and Fig. A1 B-D (constant pump).

#### 6.1.2 Modeling dynamic ion concentrations and deriving required pump densities for steady state ion concentrations

The function of the Na^+^/K^+^ pump is to maintain intra- and extracellular ion concentrations at fixed levels. If there were no Na^+^/K^+^ pump in an excitable cell, sodium ions would accumulate inside the cells and potassium ions would accumulate in extracellular space. This would reduce the concentration differences between ions in intra- and extracellular space, which impedes the firing of action potentials. The goal of the Na^+^/K^+^ pump is therefore to retain fixed sodium and potassium reversal potentials by maintaining ion homeostasis. This is achieved when the energetic supply, which would be the rearranging of ions by the Na^+^/K^+^ pump exactly equals the energetic demand which is the ion displacement caused by the action potentials.

In order to fully understand the influence of the Na^+^/K^+^ pump on cell excitability, we modeled the ion displacements of action potentials and the pump explicitly (implemented to create figures 2,4,5, and 6D). To this end, we added ion concentration dynamics of intra- and extracellular sodium and potassium to the model equations (Eqs. 7, 8, 9, and 10) similarly to (56), and introduced a dependency of the reversal potentials on these ion concentrations via the Nernst equation (eqs 11, 12):

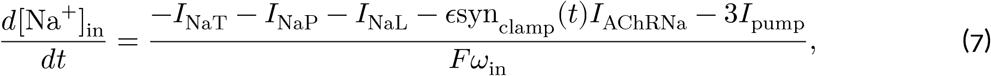

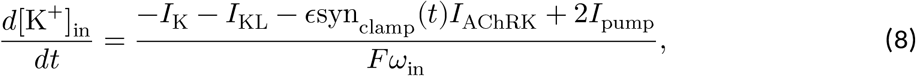

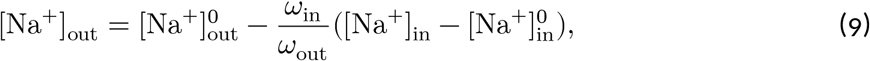

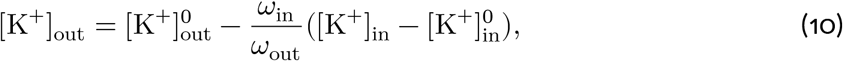

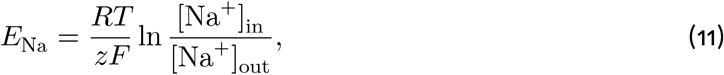

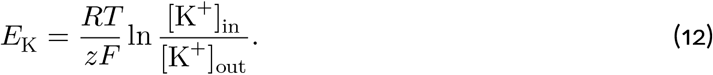

**Figure 6.**
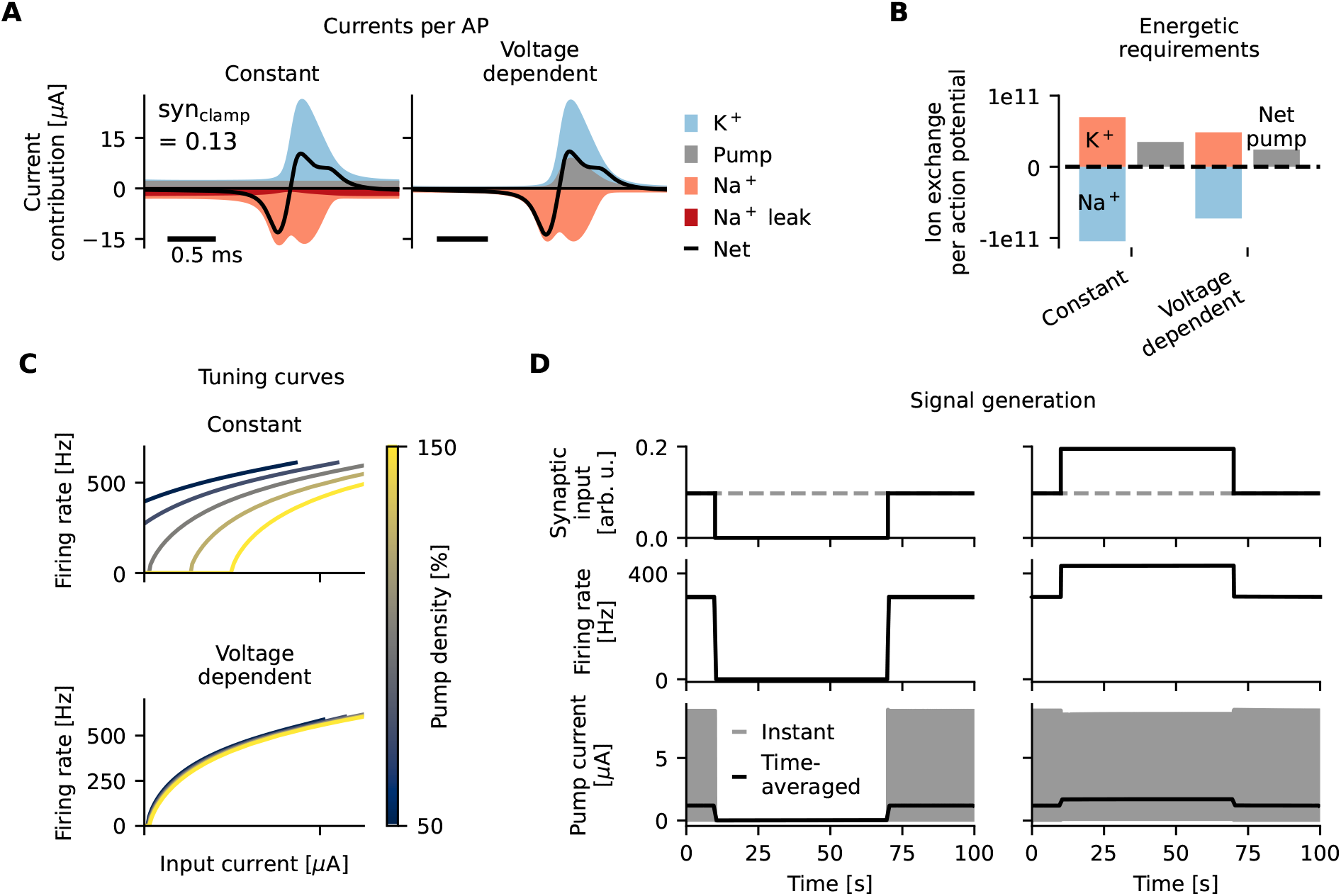
Ideal voltage dependence of the Na+/K+-ATPase for energy-efficient action potentials and
minimal firing-rate adaptation. (A) Action potential current contributions to the total in- and outward currents for a constant and physiologically relevant synaptic drive (syn_clamp_=0.13, Methods 6.2.1) with a Na^+^/K^+^-ATPase without voltage dependence (left) and with an optimal voltage dependence that mimics potassium channels (right). The voltage-dependent pump current takes on the role of a potassium channel and contributes significantly to the net current at the AP downstroke. (B) Total amount of sodium (red) and potassium (blue) ions, and net ion transfer of the pump (grey) that are relocated per AP for a cell with a voltage-insensitive pump (left) and a voltage-dependent pump (right). (C) The effect of pump density on the tuning curve is minimal for ideal voltage-dependent pumps (bottom) compared to a non-voltage-dependent pump (top). (D) Signal generation in a cell with Na^+^/K^+^-ATPases with optimal voltage dependence. Synaptic input suppression (top left) silences the cell (center left) and reduces the pump current (bottom left). Firing rates are however not changed, and the cell remains silent. Synaptic input doubling (top right) increases firing rates (center right) and increases time-averaged pump currents (bottom right). Firing rates are however not affected. Note that the instantaneous pump current (bottom, grey) varies on the timescale of action potentials, which is highly compressed in this 100 second time window.

Here, *F* is the Faraday constant and *ω*_in_ is the intracellular volume which, as measured by (49) is roughly 0.434 mm^3^. 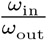 is the ratio between the volumes of the intra- and extracellular space. As electrocytes are relatively large compared to their environment, we assumed 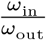 to be 2. Initial and steady-state ionic concentrations in intra- and extracellular space, 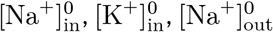, and 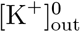, were set to 13.5 mM (see table 1), 89 mM, 120 mM, and 2.16 mM (see value reported in (26)) respectively. These values were also used to initialize simulations. Note that *I*_stim_(*t*) has been decomposed into *−ϵ*syn_clamp_(*t*)*I*_AChRNa_ and *−ϵ*syn_clamp_(*t*)*I*_AChRK_. This has been done to separately track the sodium and potassium displacement caused by the input stimulus.

To simplify the analysis, the model equations were rewritten to cancel out the state variable [K^+^]_in_, and have the model depend only on [Na^+^]_in_. To achieve this, we first rewrote the membrane potential equation, Eq. 2, so that it is separable into Na^+^ and K^+^ currents, by inserting Eqs. 4 and 1;

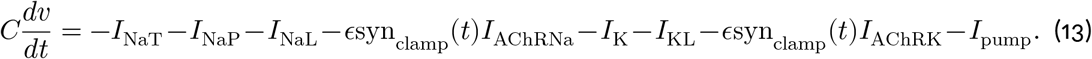

According to Eqs. 7 and 8, the membrane potential equation can thus be expressed as

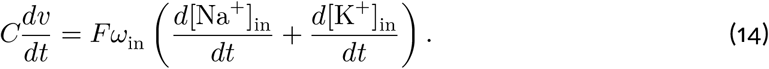

Integrating on both sides gives us

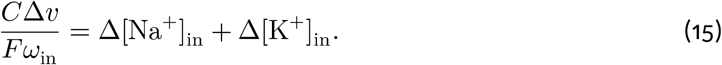

As the membrane conductance *C* is small, the Faraday constant *F* is very big, and the intracellular volume *ω*_in_ is also relatively big, we can approximate 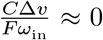. This proves that in our model, the macroscopic changes in intracellular ion concentrations can always be related by

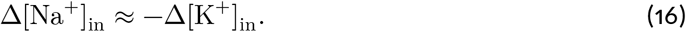

This allows us to rewrite equations 8 and 10 to depend only on state variable [Na^+^]_in_

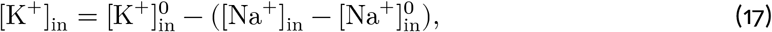

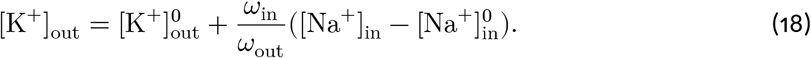

With a perfectly working Na^+^/K^+^ pump, the macroscopic change in intracellular sodium Δ[Na^+^]_in_ is zero, which signifies that the energetic supply of the pump exactly equals the energetic demand of the action potentials. From Eqs. 9, 17 and 18 we can conclude that if there is no macroscopic change in intracellular sodium, extracellular sodium concentrations and intra- and extracellular potassium concentrations also remain constant.

Under the assumption that the Na^+^/K^+^ pump is solely responsible for all active sodium transport, long-term ion homeostasis in a high frequency firing electrocyte can be obtained if we tune the baseline pump current, 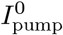, so that Δ[Na^+^]_in_ (Eq. 7) is zero i.e.

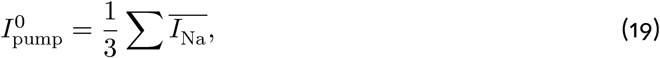

where 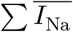 is the sum of the time average of all sodium currents, which in this model are *I*_NaT_, *I*_NaP_, *I*_NaL_, and *ϵ*syn_clamp_(*t*)*I*_AChRNa_. The time-average is here taken in a tonic firing regime where the averaging window equals one spiking period. As 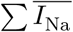 depends on 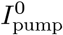 due to the co-expression of pumps and sodium leak channels (Eq. 6), we iteratively recompute 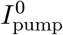 according to equation 19 until the condition is satisfied with an error margin of 1 nA. This was done to arrive at currents and APs shown in figures 1 B, H-J (with pump) and 6 A-C(constant pump). This procedure was also used to initialize simulations at steady state values in figures 2, 4, and 5. Steady state pump currents for various stimulus protocols are reported in Table 2.

**Table 2:** All baseline stimuli and tuned parameters presented in this article. 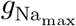 was tuned to maintain a spike amplitude of 13 mV, and 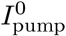 and 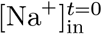 were tuned to maintain ion homeostasis (and thus steady state values) for the specified stimuli. Empty cells correspond to standard parameter values as reported in (26) and Table 1.

| Figure | Stimulus (at baseline) |  |  | Tuned parameters |  |  |
| --- | --- | --- | --- | --- | --- | --- |
| | $\text{syn}_{\text{clamp}}$<br>[arb. u.] | $r_{\text{pn}}$<br>[Hz] | $\epsilon$<br>[arb. u.] | $g_{\text{Na}_{\text{max}}}$<br>[$\mu\text{S}$] | $I_{\text{pump}}^0$<br>[$\mu\text{A}$] | $[\text{Na}^+]_{\text{in}}^{t=0}$<br>[mM] |
| 1H-J, 6A-B (left) | 0.13 | – | – | – | 1.888 | – |
| 2 | 0.9 | – | – | – | 1.630 | – |
| 4 | – | 600 | – | 1300 | 4.074 | – |
| 5B-D | – | 260 | 0.5 | 881 | 1.368 | – |
| 5E-G | – | 260 | – | 763 | 1.258 | – |
| 6 A-B (right) | 0.13 | – | – | – | – | – |
| 6D | 0.9 | – | – | – | – | 13.517 |

#### 6.1.3 Modeling the feedback loop of ion concentrations on pump density

Assuming that pump densities are tuned to sustain a fixed baseline firing rate, deviations from this baseline firing will lead to a mismatch between the ion displacement caused by action potentials and the ion restoration of the Na^+^/K^+^ pump. This will lead to a shift in intra- and extracellular ion concentrations. As the pump rate and thus *I*_pump_ is a function of intra- and extracellular ion concentrations (56), the pump rate will adjust accordingly. We model the dependency of *I*_pump_ on intracellular sodium concentrations [Na^+^]_in_ and extracellular potassium concentrations [K^+^]_out_ similarly to (56)

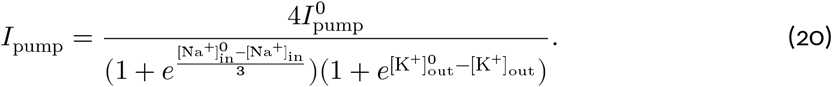

For simplicity, we adjusted the terms within the exponents so that 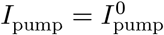 when Δ[Na^+^]_in_ = 0. Here, 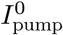 is the pump current that is tuned to facilitate ion homeostasis at the baseline firing rate. As the pump current saturates at 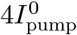, which is proportional to the number of Na^+^/K^+^-ATPases that are expressed, the baseline pump current 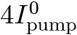 is also proportional to the pump density. A shift in *I*_pump_, which can now happen as a consequence of deviations from baseline firing, without co-expression of *g*_NaL_, which is unlikely on small timescales, will lead to a shift in cell excitability. This feedback loop is implemented and its effects shown in figures 2,4,5,and 6D.

#### 6.1.4 Mechanisms that improve entrainment

The feedback loop of ion concentrations on pump density alters pump currents when inputs change (Fig. 2), and consequently disrupts synchronization (Fig. 4, 5). Two mechanisms that alleviate these consequences of pump activity on synchronization were explored: extracellular potassium buffering and increased synaptic weights.

Extracellular potassium buffering was implemented to decrease the variation in Δ*I*_pump_, and thereby the variation in mean-driven electrocyte properties (Eq. 20, Fig. 4 D-E). Transient buffer effects were neglected, and an instantaneous potassium buffer with infinite capacity was assumed by setting

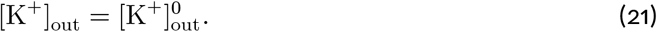

Synaptic weights were implemented by setting *ϵ* = 0.5 for weak coupling (Fig. 5 B-D), and *ϵ* = 1 for strong coupling (Eq. 1, Fig. 5 E-G).

#### 6.1.5 Modeling an optimal voltage-dependence of the pump

Dynamics of action potential firing would be unaffected by the presence of voltage-dependent electrogenic pumps if the membrane voltage would modulate their activity in a way that the pump current mimics hyperpolaring voltage-gated and leaky potassium current. We substantiate this idea by modeling a voltage-dependence of the pump that copies the dynamics of potassium currents (Fig. 6, voltage-dependent pump). To achieve this, we rewrite the baseline pump current 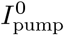 as a function of the membrane voltage, and a transformation of the membrane voltage that takes into account the history of the membrane voltage (*n*)

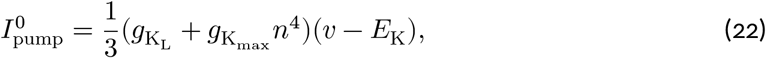

which is essentially a scaled version of a combination of the equations that describe the voltage-gated and leaky potassium currents. As the pump current now behaves like 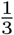 of the potassium currents, we can reduce the potassium conductances by 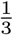 and still get qualitatively the same APs, through

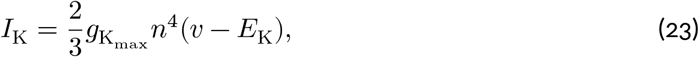

and

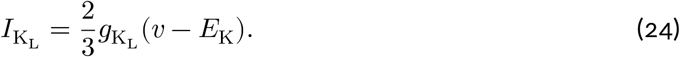

By restructuring the current and pump equations as such, the pump current should always equal ^1^ of the potassium currents, which exactly satisfies the energetic demand of the cell. There is however a third potassium current, *I*_AChRK_, that is activated by neurotransmitter release. As we cannot expect the pumps to also be sensitive to neurotransmitter release, the voltage-dependent pump described in equation 22 will pump slightly less than necessary to maintain ionic homeostasis. We therefore let the model run until steady state ion concentrations were reached, which happens in close proximity to the baseline concentration of [Na^+^]_in_ = 13.500 mM, which is [Na^+^]_in_ = 13.517 mM. We then initialize subsequent simulations with 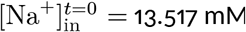.

### 6.2 Stimulus protocols

In this study, the influence of the pump current on cell excitability was studied for increasingly complex physiologically relevant stimulus protocols. For each stimulus protocol, baseline pump currents, 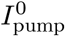, or initial intracellular sodium concentrations, 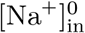, were tuned to facilitate ionic homeostasis. Furthermore, when simulating communication paradigms, voltage-gated sodium conductances, 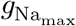, were tuned to maintain a spike amplitude of 13 mV. Stimuli and tuned parameters for each experiment shown in this article are reported in Table 2. The selected stimuli and their implementations are explained below in more detail.

#### 6.2.1 Stimulation in the mean-driven regime

For creating frequency-input curves (Fig. 1 C, F, 3 C, and 6 C), *I*_stim_(*t*) was replaced with a constant input current, and ionic concentrations were fixed to baseline values to avoid the pump-induced changes in firing rates. The *f*-*I* curves shown in this article therefore represent instantaneous firing rates. Representative inputs were estimated and used to show the influence of the pump current on tonic firing (Fig. 1 D, G). These inputs were estimated as follows: we first modeled an electrocyte with a periodic synaptic drive as in (26). The frequency of this drive was set to 400Hz, which is the mean value of reported EODfs (and thus presumable pacemaker firing rates) of 200-600Hz (29). Then, *I*_stim_(*t*) was averaged to obtain the average input current (0.63 *μ*A).

A similar approach was taken to show AP shape, current contributions, and energetic demand for a representative constant input (Fig. 1 H-J and 6 A-B). In order to add synaptic currents, which account for additional in- and outfluxes of sodium and potassium, in these experiments, the electrocyte was stimulated with a constant synaptic drive (syn_clamp_=0.13) taken from the average synaptic drive resulting from a 400Hz pacemaker stimulus.

To showcase the pump-current induced firing-rate adaptation in Fig. 2, the baseline synaptic drive was chosen as the mean syn_clamp_(*t*) for a periodic drive of 300Hz (syn_clamp_=0.09). This was chosen so that doubling the synaptic drive would result in a physiologically plausible synaptic drive of 0.18, which would be the average of a 600 Hz periodic drive.

#### 6.2.2 Periodic stimulation

Next, we showed the impact of pump-current induced firing-rate adaptation on the synchronization of an excitable cell to periodic input. In particular, we studied the entrainment of the electrocyte to the pacemaker nucleus. The pacemaker was not modeled explicitly, but only the time-dependent currents that would result from the excitatory synapse were (as in (26)). Here, the parameter that reflects the (unitless) magnitude of the synaptic conductance, syn_clamp_(*t*_pn_), is modeled by a piecewise function that resets *t*_pn_ → 0 at the pacemaker firing rate, *r*_pn_;

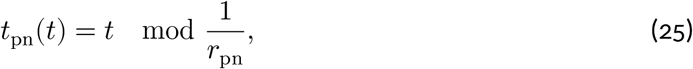

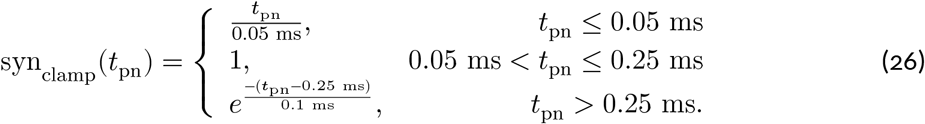

To demonstrate physiologically relevant scenarios in which a variable *r*_pn_ affects the pump current, *r*_pn_ was set to model chirps (cessations in firing) and frequency rises. To ensure a significant effect of firing-rate changes on the pump current, relatively long chirps were initiated in a fish with high baseline firing rates, and relatively high frequency rises in a fish with low baseline firing rates. Baseline firing rates, chirp duration, frequency rise amplitude, and frequency rise timescale are representative of EOD signals found in experimental settings (29).

To model chirps (Fig. 4), *r*_pn_ was set to 600 Hz, and after 100 ms, chirps were generated where *t*_pn_ was only reset after a period of 20 spikes i.e. 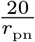. After this period, *t*_pn_ was again reset with a frequency *r*_pn_ for 100 ms. This was repeated 10 times to simulate 10 consecutive chirps.

To model frequency rises (Fig. 5), *r*_pn_ was set to 260 Hz, and after 6 seconds, frequency rises were generated where *r*_pn_ was set by the following formula

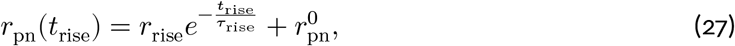

where *r*_rise_ is the amplitude of the frequency rise, which was set to 40 Hz, *t*_rise_ is the elapsed time since the onset of the frequency rise, *τ*_rise_ is the timescale of the frequency rise which was set to 1 second, and 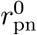 is the baseline frequency which was set to 260 Hz. Frequency rises were initiated every 2 seconds, 10 times in a row.

##### 6.2.2.1 Spike entrainment measure

To quantify the accuracy of entrainment between spikes em-mited by PN at times, 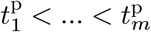 and spikes emmited by electrocyte at times 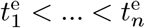, where 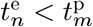.

Let 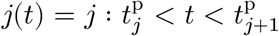 denote the index of the PN spike interval in which a given electrocyte spike time resides. Then define the phases of electrocyte spikes relative to pacemaker inter spike intervals as

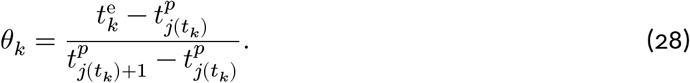

The rigidity of the entrainment phase is then quantified by the circular variance as the mean resultant length

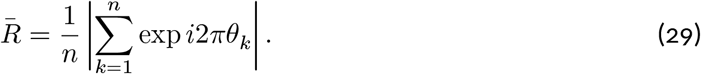

This score, which is referred to as the entrainment index, is shown on top of spike trains in figures 5 B-G.

## 7 Data availability statement

All results presented in this article can be reproduced via the code published at https://itbgit.biologie.hu-berlin.de/compneurophys_pub/electrocyte_nakatpase.

## 8 Acknowledgments

We thank Dr. Louisiane Lemaire and Mahraz Behbood for fruitful discussions. This project has received funding from the Einstein Foundation Berlin (grant number EP-2021-621).

## 9 Author contributions

Conceptualization, L.W., J-H.S and S.S.; Methodology, L.W. and J-H.S.; Software, L.W.; Writing – Original Draft, L.W., J-H.S., S.S.; Writing, – Review & Editing, L.W., J-H.S. and S.S.; Supervision, J-H.S. and S.S.; Funding Acquisition, S.S.

## 11 Appendix

## 11.1 Main results are independent from synaptic drive

As previously shown in (26), AP amplitude decreases with increasing input currents (Fig. A1 A, left). This effect remains upon addition of either a pump with constant pump rate and co-expressed sodium leak channels (Fig. A1 A, center), or a voltage-dependent pump (Fig. A1 A, right). Interestingly, even though the shape of the current contributions (Fig. A1 B) and the APs (Fig. A1 C) look very different for low (Fig. A1 C, top) and high (Fig. A1 C, bottom) inputs, the total sodium and potassium displacement per AP, and thus the pump rate, is roughly the same (Fig. A1 D). Under the assumption that voltage-gated sodium channel (NaV) expression is adjusted to facilitate fixed AP amplitudes however (as in (26)), more NaV channels would be expressed in fish with higher synaptic drives. This would then result in additional sodium influx per AP and result in higher energetic requirements per AP for electrocytes with higher firing rates (as also shown in (26)).

**Figure A1:**
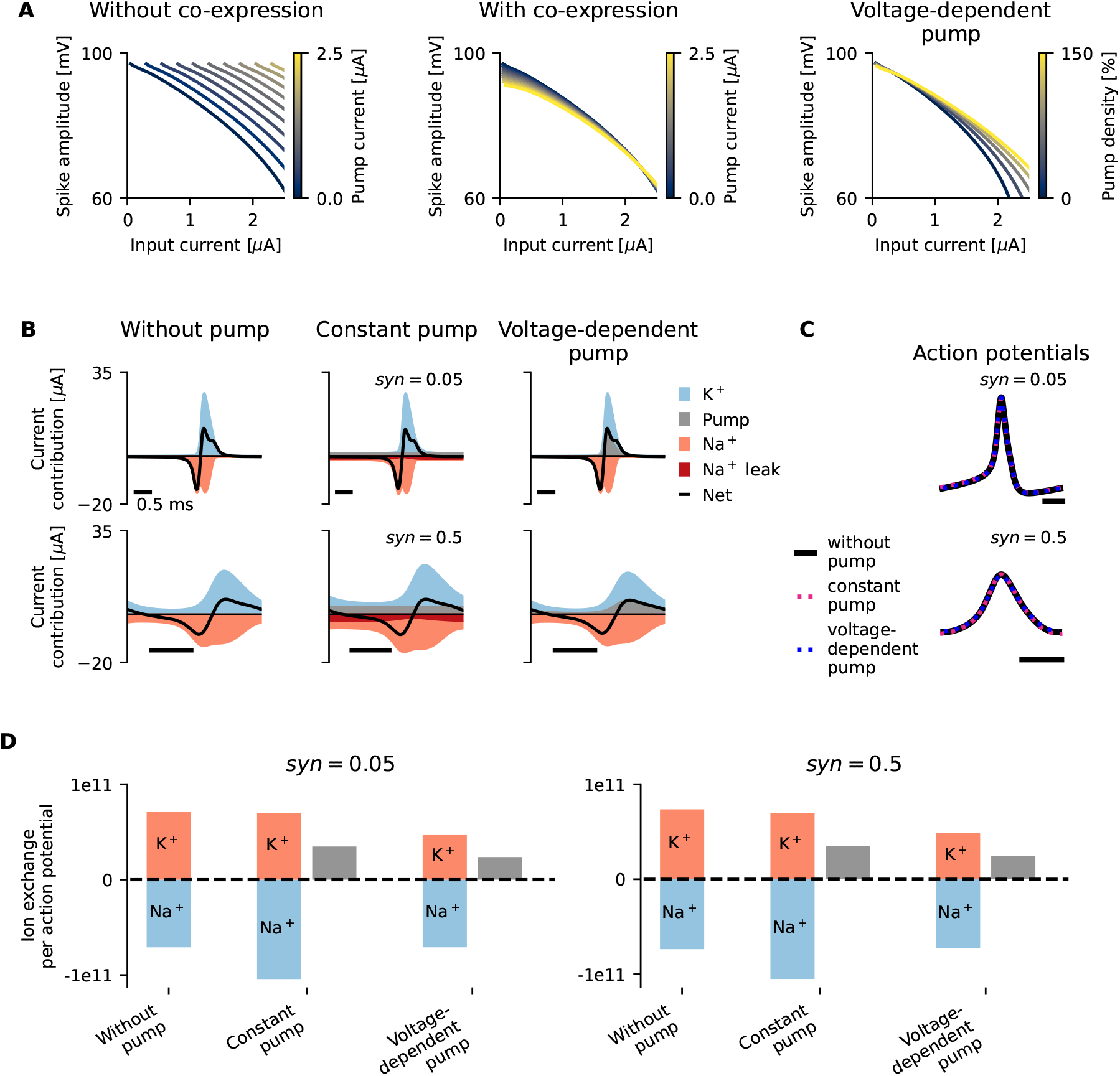
Spike amplitudes decrease with increasing input current, but current contributions per action potential (AP) remain the same. (A) Spike amplitudes decrease with increasing input currents. Without any compensatory mechanisms (i.e. co-expression of sodium leak channels, left), the pump current shifts the influence of input current on spike amplitude similarly to that on the frequency-vs-input curve (Fig. 1 C). Pump currents that are regulated either through co-expressed channels (center) or a voltage-dependence of the pump (right) have little influence on the relation between input current and spike amplitude. (B) A comparison of all currents that contribute to the AP between constant small (top, syn=0.05), and large (bottom, syn=0.5) synaptic inputs which result in low and high firing rates respectively. (C) AP shapes of electrocyte models without and with a constant or voltage-dependent pump current are indistinguishable. (D) Ion exchanges per AP, and thus pump rates, are similar for low (left) and high (right) synaptic inputs.

## 11.2 Phase oscillator theory to quantify entrainment

The observation that each PN spike typically causes one electrocyte spike suggests that electrocytes are excitable cells, kicked above threshold by synaptic PN inputs and then return to rest. However, taking into account the sustained high frequency PN firing rates, *r*_pn_, and the kinetics of the electrocyte acetylcholine receptor shows that the model is better understood as a mean-driven entrained oscillator. To see this, the membrane equation of the electrocyte, Eq. 2, is rewritten to separate the stimulus current into a time-averaged DC component, Eq. 30, and a time-dependent zero-mean component.

Averaging the stimulus over one PN period gives

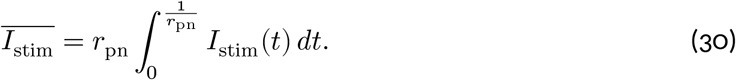

As will be shown below, in many cases, the mean drive, 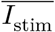, is sufficient to induce spiking the the electrocyte. Reformulating Eq. 2 to include the mean stimulus expressed in Eq. 30 reads

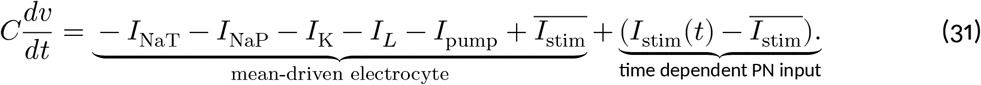

As the baseline pump current, which depends on the mean input (see Methods 6.1.2), is compensated for through the co-expression of sodium leak channels to generate similar mean-driven dynamics for different baseline pump currents (see Fig. 1 F and Methods 6.1.1), the baseline pump current, *I*_pump_, is here set to zero to simplify analysis. Note that mean-driven properties will be qualitatively similar, but slightly different depending on the baseline pump current, and thus the co-expressed sodium leak channels (as also shown in slightly varying fI curves in Fig. 1 F).

We can now characterize the mean-driven electrocyte by the relation of its firing rate to mean input in terms of its *f*-*I* curve (Fig. 3 E). We can furthermore determine the time averaged stimulus for various PN driving frequencies (Fig. 3 F). When comparing the mean-driven frequency of the electrocyte, *r*_*e*_, to the PN driving frequency, *r*_pn_, we find mismatches in frequencies that can go up to 180 Hz (Fig. 3 G). As proven by (26), the electrocyte model can be entrained by *r*_pn_ ranging from 200 to 600 Hz. We can therefore assume that the input stimulus is strong enough to overcome large frequency mismatches between *r*_*e*_ and *r*_pn_.

To quantify the allowed frequency mismatches between *r*_*e*_ and *r*_pn_ for synchronization, we treat the mean-driven electrocyte as a periodic oscillator that is to be entrained by an external force, which is the PN. The evolution of the phase of the electrocyte can now be expressed as

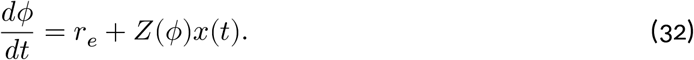

Here, *r*_*e*_ is the frequency of the mean-driven electrocyte, *Z*(*ϕ*) is the change in phase of the electrocyte evoked by perturbations of the membrane potential, and *x*(*t*) is the time-dependent perturbation caused by the zero mean pacemaker input stimulus

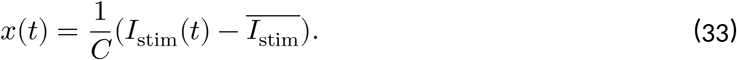

One can now define a variable that describes the phase difference between the electrocyte and the PN as

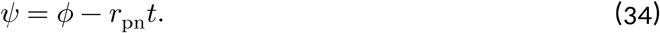

If there exists a phase *ψ* for which *ψ* does not change over time i.e.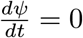, the electrocyte and the PN will be phase-locked, or entrained. The equation for evolution of the phase difference is obtained by plugging Eq. 34 in Eq. 32

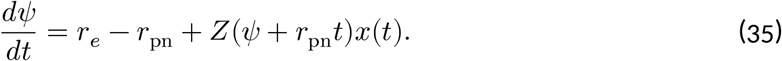

As *ψ* ≪ *r*_pn_*t* is a slow variable, one period of *ψ* will ‘see’ a lot of PN periods. Hence, the method of averaging yields

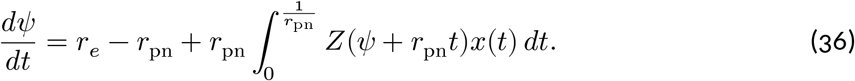

From this equations the minimum and maximum *r*_pn_ for which the electrocyte and the PN can be phase-locked is defined as

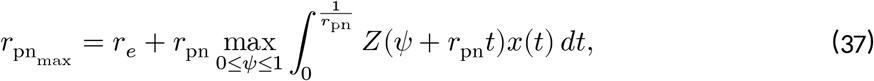

and

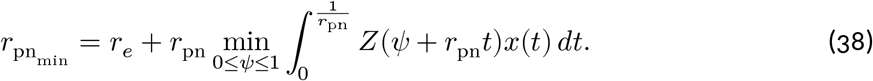

The range 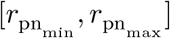 is referred to as the entrainment range.

The mean-driven features *r*_*e*_ and *Z*(*ϕ*) are altered upon a deviation of the pump current Δ*I*_pump_ (Fig. A2 A, B), which is in essence the same as a deviation in 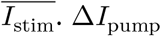 thus affects 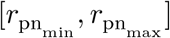 and thereby the entrainment of the pacemaker-driven electrocyte (Fig. A2 C).

**Figure A2:**
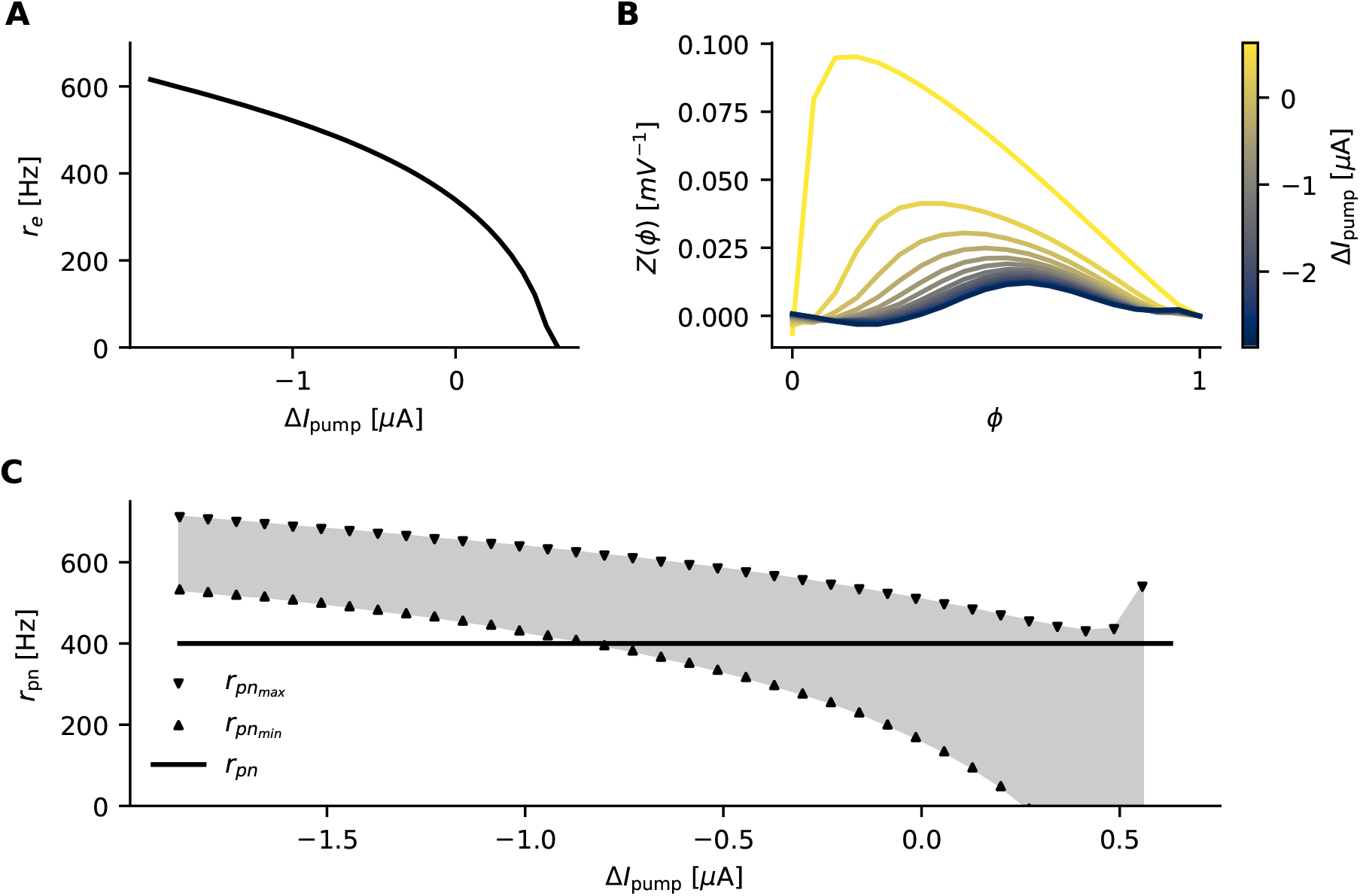
A deviation in pump current ΔI_pump_ alters mean-driven electrocyte properties and thereby its entrainment region. (A) Mean-driven electrocyte frequency r_e_ as a function of ΔI_pump_. (B) Phase Response Curves (PRCs, Z(ϕ)) as a function of ΔI_pump_. (C) The entrainment range 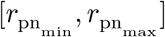, which is a function of mean-driven electrocyte properties (A, B, Eqs. 38, 37), changes upon deviations in pump current. For very strong deviations in I_pump_, the pacemaker frequency r_pn_ falls out of the entrainment range which means that the electrocyte will not lock to the pacemaker in this regime.

The influence of the synaptic weight on entrainment (Fig. 5 C-H) can also be explained by equations 38, 37, as it increases x(t) and therefore 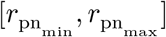.

### 11.2.1 Phase Response Curves

To solve Eqs. 37 and 38, PRCs need to be computed. PRCs were obtained for various 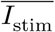 by first injecting constant input current 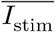 to the electrocyte. Then, the phase of the electrocyte was defined as the peak of one spike in the tonically firing electrocyte to the next peak. The membrane voltage was then perturbed by 1mV at 20 different phases linearly interpolated between 0 and 1, and the resulting time delays or advances of the next spike was recorded as the phase response.

## 11.3 Relation between integrated sodium and potassium currents over one ISI

In a conductance-based model without pump, where all currents are sodium or potassium based (which is the case in the model used for this study, as shown in Eq. 13, if the pump current is set to zero), the cumulative sodium and potassium currents have to add up to zero between two spikes. This becomes evident when expressing the membrane equation as such;

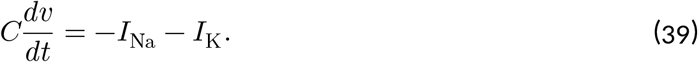

If *v* is in a limit cycle with length *T*,

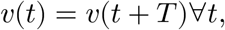

and

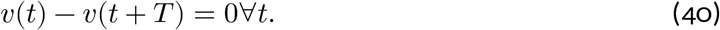

If we integrate the voltage equation (Eq. 39) on both sides from *t* to *t* + *T*, we get

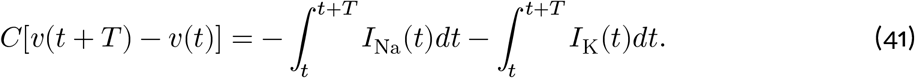

If we plug in Eq. 40, we get

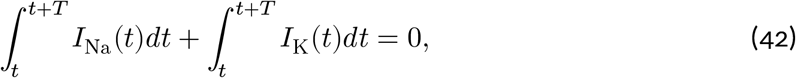

which means that the integrated sodium and potassium currents over the time course of one ISI should always be equal to each other.

## 11.4 Considerations on metabolic costs of compensatory mechanisms

Quantitative estimates of metabolic costs in this study are based on the ATP that is required to fuel the Na^+^/K^+^ pump. This includes the cost of the restoration of sodium and potassium ions that flow to support action potentials, resting potentials, and postsynaptic potentials.

The co-expression of pumps and sodium leak channels (see Fig. 1) and even an ideal voltage dependence of the pump (see Fig. 6) have a direct impact on the metabolic cost related to this ATP-fueled Na^+^/K^+^ pump. By integrating the net pump current over time and dividing by one elemental charge, we find the rate of ATP that is consumed for either compensatory mechanism. When compensating a relatively ‘constant’ Na^+^/K^+^-pump current with sodium leak channels, the amount of ATP spent on pumping sodium is 33% higher than it would be for a voltage-dependent pump (see Methods 6.1.5).

The impact that either of these compensatory mechanisms have on the whole cell, however, also depends on other costs, such as those related to cellular maintenance. A voltage-dependent pump would save costs related to Na^+^/K^+^ pumping, which, based on energy budgets formerly estimated for AP-firing neurons in the brain (96), is likely to be one of the main contributors to the total metabolic cost (in cerebellar cortex, for example, amounting to >50% of the total metabolic cost). Because the peak load of a voltage-dependent pump, however, is four times higher than a relatively constant pump, four times more Na^+^/K^+^ pumps would need to be expressed on the cell membrane. To be more exact, if a single pump translocates around 450 sodium ions per second (97), 8×10^10^ pumps are required to support the constant pumps and 32×10^10^ pumps are needed to support voltage-dependent pumping. If one assumes the electrocyte is a perfect cylinder, and its membrane surface were smooth (an approximation not too realistic), the total available membrane space would be 3.4 mm^2^ (49). If the Na^+^/K^+^ ATPase expression density would be as high as in the outer medulla of rabbit kidney (98), where ATPases are densely packed, a smooth electrocyte membrane would ‘fit’ 4.2×10^10^ pumps, which is two times less than necessary for constant pumping, and eight times less than required for voltage-dependent pumps. According to our model, therefore, the invaginations on the posterior side of the membrane (49) are necessary to drastically increase membrane area in order to support the large number of pumps required for ion restoration. This, in turn, would increase the ‘housekeeping’ costs of the cell related to turnover of macromolecules, axoplasmic transport, and mitochondrial proton leak, which in different brain areas are estimated to occupy 25-50% of the total energy budget (30,99). As there is insufficient data on the ratio between costs related to Na^+^/K^+^ pumping and ‘housekeeping costs’, and the fraction of housekeeping costs related to Na^+^/K^+^-pump maintenance, a quantitative comparison of the metabolic cost of the two compensatory mechanisms remains challenging. Future experiments that would aid in answering this question could involve blockage of electrocyte Na^+^/K^+^ pumps and comparing oxygen consumption to a control where electrocyte Na^+^/K^+^ pumps are functional.

Another compensatory mechanism that was discussed in this article is extracellular potassium buffering (see Fig. 4), which in electrocytes likely occurs via its extensive capillary beds (26,49) that transport excess extracellular potassium to the kidney. Assuming that an equal amount of ATP is needed in total to fuel Na^+^/K^+^ pumps, either all in the electrocyte, or partly at the electrocyte and partly in the kidney, the additional costs incurred by the extracellular potassium buffer would be dominated by the structural and maintenance costs of the capillaries. We are, however, not aware of an accurate estimate of these costs, especially since the capillaries also have additional functions such as providing other resources and transporting other waste products.

Lastly, a strong synapse was said in the article to support cell entrainment under fluctuating pump currents (see Fig. 5), but also to incur additional metabolic costs. In the example shown in the main text, however, baseline Na^+^/K^+^ costs are smaller for a stronger synapse, see Fig. 5B (weak synapse) vs. Fig. 5E (strong synapse). This is the case because, similarly as shown in (26) Fig. 7B, a weak synapse elicits smaller postsynaptic potentials, which lowers the AP peak with respect to a stronger synapse. To make a fair comparison on the metabolic costs between a weak and a strong synapse, voltage-gated sodium conductances were scaled to maintain a peak amplitude of 13 mV (see Methods 6.2). For weak synaptic stimulation, a higher voltage-gated sodium conductance was needed to reach this peak amplitude, which, due to the excess inflow of sodium through these voltage-gated channels resulted in an increase of 10% in ATP consumption by Na^+^/K^+^ pumps with respect to strong synaptic stimulation.

There are, however, additional costs that scale with synapse strength, such as the restoration of presynaptic calcium, the restoration of (presumably small amounts of) postsynaptic calcium, and neurotransmitter packaging, and recycling. In the brain, these costs are estimated to be 0.18 to 1 times the cost of fueling the Na^+^/K^+^ pumps that restore the sodium ions that traverse neurotransmitter receptor channels (96,100). In our model, merely 11% of sodium ions enter the electrocyte via neurotransmitter receptor channels in the strong-synapse case. Assuming that above-mentioned additional costs are equal to those related to Na^+^/K^+^ pumping of neurotransmitter-related currents (according to the upper bound estimate by (100)), a weak synapse (half the size of the strong synapse) would incur a cost increase of 5.5% and a strong synapse would incur an increase of 11%. This would, however, still result in a 4% higher cost efficiency of a strong synapse compared to a weak synapse.

There is reason to believe that the fraction of the energy budget related to the restoration of presynaptic calcium, the restoration of (presumably small amounts of) postsynaptic calcium, and neurotransmitter packaging, and recycling in the electrocyte could differ significantly from those estimated by (96,100). First, to the best of our knowledge, such energy budget estimations have only been done for neurons active at significantly lower firing rates than electroyctes (by a factor of approximately 100), and, second, operate mostly under the glutamate neurotransmitter, while electrocyte receptor channels are activated by acetylcholine. An accurate estimate of the impact of synapse strength on the electrocyte energy budget, therefore, requires quantitative data on the rapid dynamics of acetylcholine production in the presynaptic neuron and recycling in the synaptic cleft, which, currently is also hard to estimate.

Supported by the above-mentioned considerations, we argue that the impact of mechanisms that compensate for Na^+^/K^+^-pump currents on an electrocyte’s metabolic cost could be significant. Due to the absence of more detailed experimental quantification, a plausible quantitative cost estimate remains beyond the scope of this article. We note, however, that although the metabolic costs of potassium buffering and synaptic strength is likely to differ between cell types, the energetic estimate of the respective ATP requirements by Na^+^/K^+^ pumps for constant vs. voltage-dependent pumping generalizes and extends to all excitable cell types (as is discussed in the main text, section 5.1).

